# Attention reshapes the information dynamics of thalamic and cortical prediction-error learning

**DOI:** 10.64898/2026.07.10.737794

**Authors:** Juho Äijälä, Michael A. Jensen, Kai J. Miller, Dora Hermes, Alejandro O. Blenkmann, Robin A.A. Ince, Tristan A. Bekinschtein, Martin Vinck, Max Garagnani, Andres Canales-Johnson

## Abstract

Prediction errors (PEs) drive perceptual learning by updating internal models of the sensory environment, yet it remains unclear how attention reshapes their representation across distributed thalamocortical circuits. Using intracranial stereoelectroencephalography (sEEG) from 17 patients performing a roving auditory oddball task under attended and unattended conditions, we quantified PE encoding using mutual information and co-information to capture redundant and synergistic PE representations. Attention modulated PE encoding in both the thalamus and the temporal cortex, but with distinct informational dynamics. Thalamic encoding showed a stable reduction of PE information during distraction, consistent with state-dependent thalamocortical gating. In contrast, the temporal cortex expressed two opposing learning trajectories during attended listening that converged once attention was diverted, revealing distinct cortical learning regimes rather than a uniform attentional effect. Attention further reorganized the informational content of cortical PE representations by altering the balance between redundant and synergistic information. A biologically constrained neural network showed that attention-dependent changes in inhibition and long-range connectivity reproduced these dynamics through Hebbian learning. Together, these findings suggest that attention regulates predictive learning not simply by changing the strength of PE responses, but by reshaping how distributed thalamocortical circuits represent and integrate sensory evidence over time.

## Introduction

Theories of perception propose that the brain continuously compares top-down predictions with incoming sensory signals and uses prediction errors (PEs) to update internal models of the environment ^1–4^. In the auditory system, such PE responses have been observed across various scales, species, and states of consciousness and are classically indexed by event-related potentials (ERPs) ^3,5–8^. Converging evidence from intracranial and large-scale electrophysiological studies suggests that PE signals are often distributed across multiple brain regions rather than localized to a single cortical source ^5,9–13^. This shifts the central question from whether PEs are present to how they are represented across circuits, and how those representations are reshaped by behavioural state.

A key implication of distributed PE coding is that its informational architecture may differ qualitatively across circuits. Signals can carry overlapping information about a prediction error, yielding redundancy, or complementary information that becomes available only when signals are considered jointly, yielding synergy ^14,15^. This distinction is especially relevant for predictive processing, where PE computation is not merely local mismatch detection but rather part of a recurrent hierarchical process that selects, routes, and integrates evidence to update internal models ^3,4^. Recent work has shown that distributed cortical PE representations can be highly synergistic, indicating that a substantial fraction of PE information resides in interactions among signals rather than in any single response ^14^. Because recurrent predictive computations are thought to recruit both corticocortical and thalamocortical loops, these findings motivate a thalamocortical account of PE coding in which sub-cortical and cortical signals jointly contribute to the representational structure of prediction errors.

Attention provides a particularly important test case for this question. In predictive-processing accounts, attention is often formalized as precision weighting: a context-dependent modulation of the gain assigned to PE signals, determining whether mismatches are amplified to drive updating or down-weighted as noise ^16–18^. Mechanistically, such precision control has been linked to changes in gain and effective connectivity within predictive circuits ^2,19–21^, implying that attention could reshape not only the magnitude of mismatch responses but also the informational architecture of PE coding across distributed circuits. Crucially, because prediction errors are evaluated against learned sensory regularities, attentional effects on PE signals may depend not only on current task state but also on how expectations are formed and revised over repeated stimulus exposure ^22–25^. This possibility is especially relevant in paradigms such as the roving oddball, where repeated standards progressively establish auditory regularities and shape subsequent mismatch responses. The literature on attention and mismatch responses has nevertheless been notably mixed: some studies report enhancement of PE-related responses under attention, others attenuation under competing task demands, and others little group-level change despite substantial individual variability ^19,20,26–29^.

One possible reason for this inconsistency is that attention may not exert a uniform effect on PE responses but instead interacts with ongoing learning dynamics differently across levels of the thalamocortical hierarchy. This possibility is reinforced by growing evidence that thalamic circuitry contributes to sensory deviance detection, gain regulation, and the prioritization of PE signals, positioning the thalamus as a candidate locus for relatively stable state-dependent control over PE coding ^30^.

In fact, recent human intracranial work has shown that auditory oddball learning in the hippocampus can occur even in the absence of consciousness ^31^. Neuropixel recordings from anaesthetised patients revealed prediction error responses in both single-unit and local field potential activity, and showed that this encoding increased over the course of the experiment, which suggests representational PE-learning during unconsciousness in sub-cortical hippocampal areas. Together with research showing synergistic encoding of prediction errors across the cortical hierarchy ^14^, these findings raise a complementary question: are PE information and learning dynamics in the thalamic and cortical circuits modulated by attention? If so, what are the underlying neural mechanisms?

Here, we addressed this question by recording sEEG from the temporal neocortex and thalamus during a roving auditory oddball task performed under attended and unattended conditions. Using sEEG in patients with temporal cortical and thalamic coverage, we quantified time-resolved PE information using mutual information and decomposed temporal response information into redundant and synergistic components across pairs of time points during a roving auditory oddball task performed either while attending to tones or while engaged in a competing visual task. We hypothesized that distraction would produce a relatively stable shift in thalamic PE information, consistent with a broader state-dependent modulation of thalamocortical processing, whereas temporal-cortical PE information would show stronger dependence on repetition-dependent learning and greater inter-individual heterogeneity across the course of the experiment. Finally, because the empirical data revealed pronounced subject-level diversity, we used a brain-constrained neurocomputational model implementing Hebbian learning to reproduce and explain, at the level of cortical circuits, the observed temporal trajectories and their associated informational signatures, which the model suggests may be emerging as a result of the interactive effects between attention and learning over time.

## Results

We analysed intracranial sEEG recordings from 17 patients with thalamic and temporal coverage during a roving auditory oddball task performed under two conditions: *Attended*, in which patients listened to the tones, and *Unattended*, in which they performed a competing visual task while the tones played in the background. All patients completed the tasks in the same order, with Attended always preceding Unattended.

Before assessing neural effects, we verified participants’ engagement with the visual counting task used in the Unattended condition. During this task, patients viewed images containing numbers and counted the images containing the target number while the auditory oddball sequence continued in the background. The behavioural score was defined as the proportion of counted target-number images out of all presented target-number images. Numeric count scores were available for 16 of the 17 patients. For one patient, intense nystagmus prevented collection of a numeric count score. Across these 16 patients, mean visual-task performance was 95.4% correct (s.d. = 7.5%; range = 80–100%), indicating reliable engagement with the visual task and supporting the intended diversion of attention away from the auditory stream. In the Attended condition, in contrast, patients were simply instructed to focus on the auditory stimuli while staring at a blank screen, and no behavioural response was collected.

### Attention modulates thalamic and temporal PE information

We first asked whether attention changes the amount of PE information present in thalamic and temporal-cortex responses. To quantify PE encoding, we computed the mutual information (MI; in bits) between the neural responses and stimulus identity (i.e., whether a tone was a standard or a deviant). The MI thus indicates to what extent neural responses distinguish standards from deviants. MI was estimated using the Gaussian-copula framework ^32^ previously applied to prediction-error responses ^14^. If attention acted uniformly across the thalamocortical system, we would expect a broadly similar direction of MI modulation across regions and participants. If, instead, attention interacted differently with subcortical and cortical PE coding, MI should reveal distinct regional or subject-level patterns.

MI was computed for every electrode in each region and condition (Attended: *n* = 120 trials; Unattended: *n* = 120 trials). Condition differences were assessed within electrodes using non-parametric permutation testing (*n* = 1000; max-statistic correction across time points within each electrode; FWER = 0.05), and electrodes showing significant MI modulation were retained for subsequent analyses and visualization. Thus, the reported electrode counts summarise within-electrode corrected effects rather than a single across-electrode family-wise test. Fig. 2 illustrates the electrodes with the highest MI modulation for four example patients, showing thalamic and temporal electrodes together with ERP waveforms and corresponding MI time courses for both conditions. Fig. 3 summarises the empirical MI modulation and Bayesian prevalence patterns across regions, with thalamic results in panels **a–c** and temporal results in panels **d–i**.

**Figure 1:**
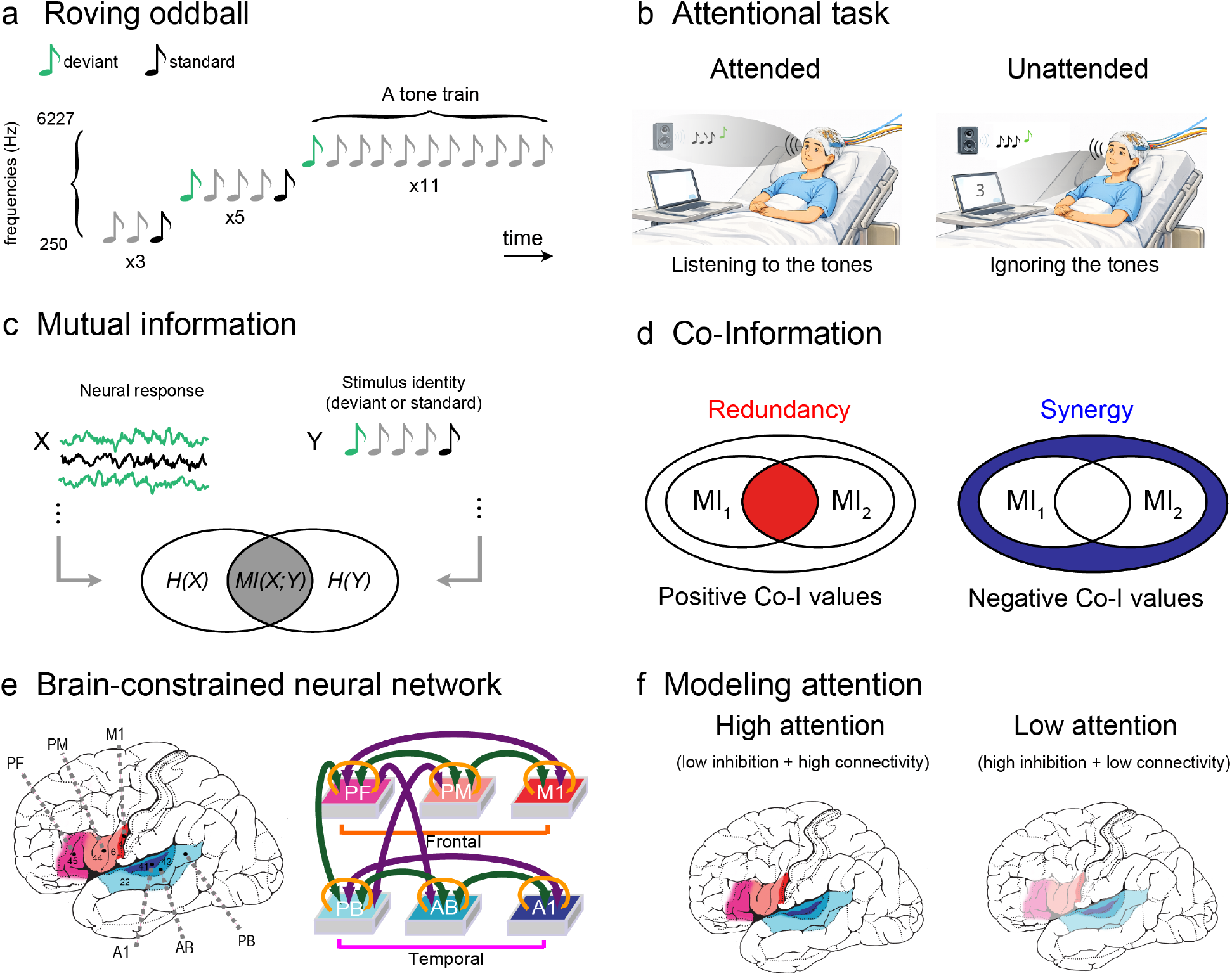
Experimental and analytical framework. **a**, Roving auditory oddball sequence. Each tone train contained repeated tones of the same frequency; the first tone after a frequency change was treated as a deviant, and later repetitions within the same train were treated as standards. **b**, Attentional manipulation. In the Attended condition, patients listened to the tones; in the Unattended condition, the auditory sequence continued while patients performed a competing visual counting task. **c**, Mutual information analysis. Trial-wise sEEG responses to deviant and standard tones were used to estimate the amount of information the neural responses carried about stimulus class. **d**, Co-information analysis. Positive CoI values indicate redundancy, where two response time points carry overlapping stimulus information; negative CoI values indicate synergy, where information is available from the joint pattern across time points. **e**, Brain-constrained neural-network model. The model contained reciprocally connected frontal and temporal cortical areas, including temporal auditory areas A1, AB, and PB used for the simulated neural readout. **f**, Modelled attention. High and low attention were implemented as paired regimes of global inhibition (GI) and between-area connectivity (Ffb), testing whether attention-like control could reproduce the empirical temporal dynamics.

**Figure 2:**
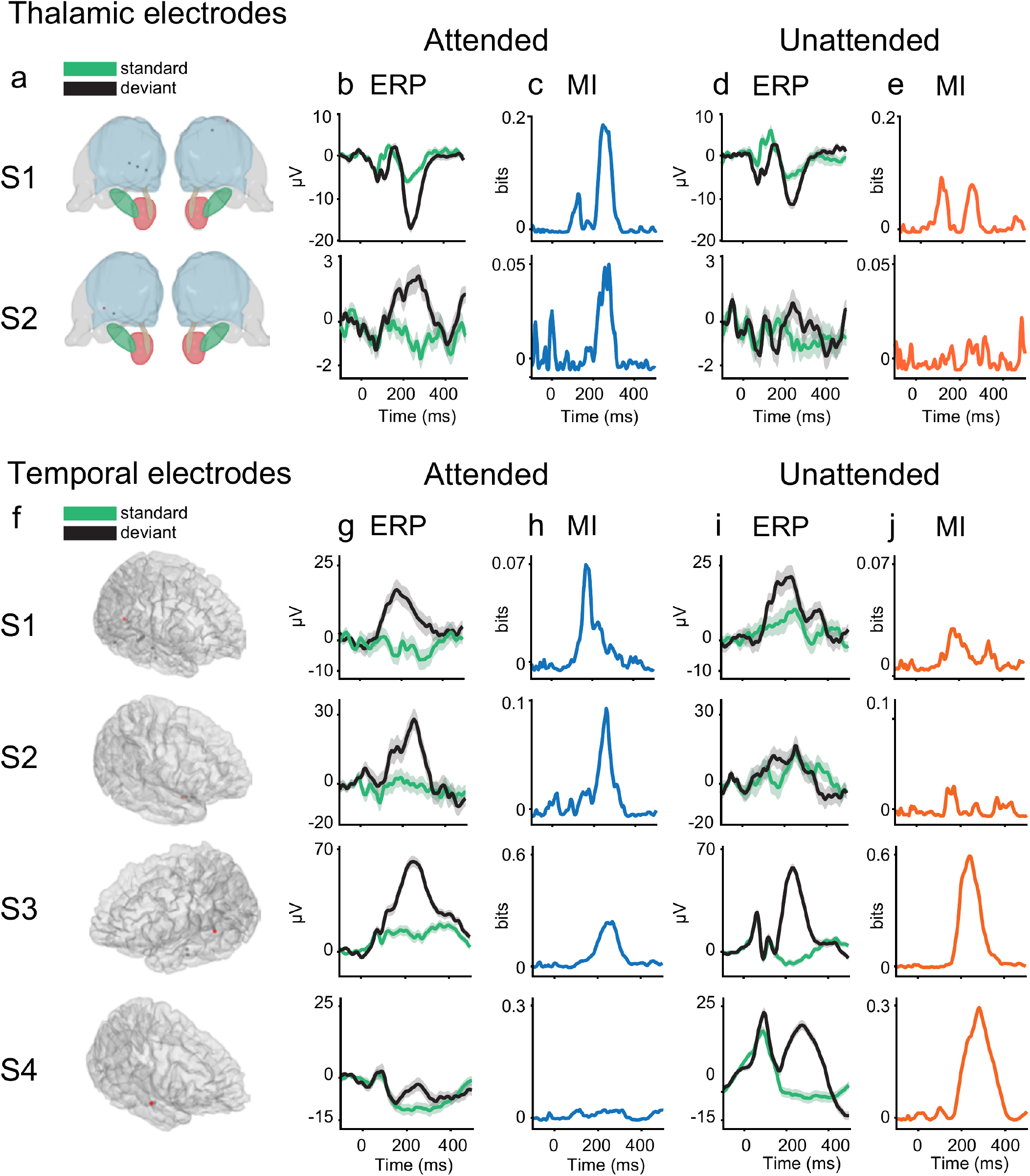
Representative thalamic and temporal modulated electrodes. Thalamic modulated electrodes for S1 and S2 (**a**) and temporal modulated electrodes for S1–S4 (**f**) are shown on 3D reconstructions. In the thalamic renderings, blue denotes the thalamic body; dark green, pink, and gold denote the subthalamic nucleus, red nucleus, and mammillothalamic tract. The red electrode marks the highest-modulated electrode in the displayed set and is the electrode used for the ERP and MI traces; grey electrodes indicate the other modulated electrodes. ERP waveforms for standard (STD, green) and deviant (DEV, black) stimuli are shown for the Attended (**b,g**) and Unattended (**d,i**) conditions, with shaded envelopes indicating SEM. Corresponding MI time courses are shown for the Attended (**c,h**) and Unattended (**e,j**) conditions. Here, MI quantifies the statistical dependence between stimulus identity and the sEEG response over time.

**Figure 3:**
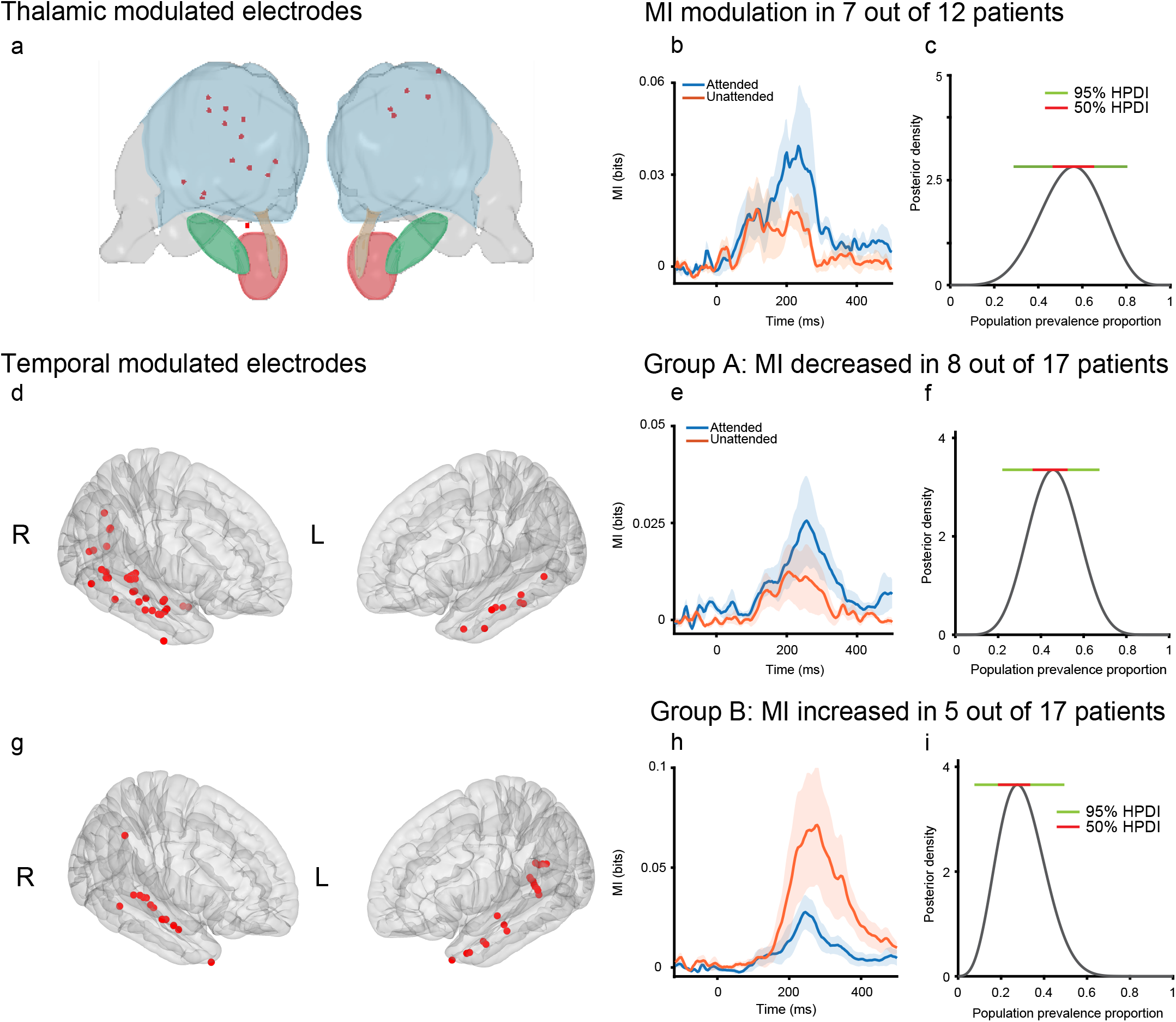
Empirical MI modulation in thalamic and temporal electrodes. (**a**) Thalamic electrodes showing significant attention-related MI modulation, displayed on bilateral thalamic renderings. In the thalamic renderings, blue denotes the thalamic body; dark green, pink, and gold denote the subthalamic nucleus, red nucleus, and mammillothalamic tract. (**b**) Group-average thalamic MI time courses for the 7 of 12 patients with significant thalamic modulation, shown separately for Attended (blue) and Unattended (orange) conditions. (**c**) Bayesian posterior distribution for the prevalence of thalamic MI modulation among comparable patients under the same experimental and analytical procedure; green and red horizontal bars show the 95% and 50% HPDIs, respectively. (**d**) Temporal electrodes in Group A, defined by lower temporal MI in Unattended than Attended. (**e**) Group A temporal MI time courses (8/17 patients). (**f**) Bayesian prevalence posterior for the Group A pattern, estimated using the full patient sample as the denominator. (**g**) Temporal electrodes in Group B, defined by higher temporal MI in Unattended than Attended. (**h**) Group B temporal MI time courses (5/17 patients). (**i**) Bayesian prevalence posterior for the Group B pattern, estimated using the full patient sample as the denominator. Shaded bands around MI traces indicate ± SEM across participants; time is relative to stimulus onset.

We first asked whether the thalamus showed attentional modulation of PEs. We found that, in total, 7/12 patients with thalamic electrodes showed significant MI modulation in at least one thalamic electrode (Fig. 3**a–c**). Of these, 6/7 showed higher average thalamic MI in the Attended condition relative to the Unattended condition, whereas 1/7 showed the opposite pattern. At the electrode level, 19/86 thalamic contacts (22.1%) showed significant MI modulation. Among significant thalamic contacts, the dominant pattern was Attended > Unattended MI (16/19, 84%), rather than Unattended > Attended MI (3/19, 16%).

Significant thalamic electrodes were concentrated in ventrolateral motor-thalamic area (Fig. 3**a**). Of the 19 significant thalamic electrodes, 15 were localized to the ventral anterior/ventrolateral border region (Thal-Va-Vl), two to the ventrolateral nucleus (Thal-Vl), one to the dorsal subdivision of the posterior ventrolateral nucleus (Thal-Vlpd), and one to a centromedian/ventrolateral/ventral posterior border region (Thal-CM-Vl-Vp). Overall, all significant thalamic electrodes involved ventrolateral (VL) thalamic territory.

To estimate the prevalence of this effect among comparable patients under the same experimental and analytical procedures, we used Bayesian prevalence ^33,34^. Given that *x* out of *y* patients show an above-chance effect, this approach estimates the expected prevalence of that effect in such patients. For thalamic MI modulation, the posterior prevalence estimate was posterior median = 0.553, MAP = 0.561, with a 95% HPDI of [0.288, 0.805] (Fig. 3**c**). This indicates that the most probable prevalence of detectable thalamic MI modulation is approximately 0.56. Thus, more than half of comparable patients would be expected to show this effect, although the uncertainty interval remains broad, spanning values from 0.288 to 0.805.

We then applied the same analysis to temporal-cortex electrodes. Here, 13/17 patients showed significant MI modulation in at least one cortical electrode (Fig. 3**d,g**). At the electrode level, 78/478 cortical contacts (16.3%) showed significant MI modulation. Significant effects were concentrated in Superior temporal gyrus (29/78, 37%), Middle temporal gyrus (14/78, 18%), and Lateral superior temporal sulcus (13/78, 17%).

Among the 13/17 patients with significant temporal-cortex MI modulation, 8/13 showed higher cortical MI in the Attended condition, whereas 5/13 showed the opposite pattern, with higher cortical MI in the Unattended condition (Fig. 3**e,h**). We refer to these two subgroups as Group A and Group B, respectively. For prevalence estimation, Group A and Group B were treated as mutually exclusive outcome categories within the full temporal-cortex sample (*n* = 17). For Group A (8/17) the Bayesian prevalence estimate was posterior median = 0.445, MAP = 0.443, with a 95% HPDI of [0.219, 0.674] (Fig. 3**f**).

For Group B (5/17), the corresponding Bayesian prevalence estimate was posterior median = 0.273, MAP = 0.257, with a 95% HPDI of [0.076, 0.495] (Fig. 3**i**). Thus, among comparable patients under the same experimental and analytical procedure, the most likely prevalence of the Group A pattern is approximately 0.44, whereas the Group B pattern is expected in approximately 0.26. The 95% HPDIs indicate uncertainty around these estimates, with values ranging from 0.219 to 0.674 for Group A and from 0.076 to 0.495 for Group B.

The subgrouping of patients into two groups was not explained by electrode sampling location within the temporal cortex. In a participant-level permutation analysis, Group A and Group B did not differ in the distribution of significant temporal electrodes across subregions (*P* = 0.382). In a complementary electrode-level analysis, the direction of electrode-level modulation was also not associated with temporal subregion identity (*P* = 0.300; see Methods).

However, these anatomical summaries and tests should not be interpreted as fine spatial localization of the underlying generators. Because MI was computed from ERP-like field-potential responses, electrode-level effects reflect the spatial listening zone of intracranial field-potential recordings, which depends on frequency content, electrode geometry, and referencing scheme ^35^.

### Empirical redundancy and synergy of PE representations

After quantifying the amount of information in thalamic and temporo-cortical areas, we aimed to determine the type of information being represented about PEs. As distributed representations of PE have been shown to encode both complementary (synergistic) and shared (redundant) information ^14^, we asked how attention can modulate this information representation in cortical and subcortical structures.

To this end, we employed co-Information (co-I), an information-theoretic measure that quantifies the balance of redundant and synergistic information in the signal. Positive co-I values indicate redundancy: two time points carry overlapping information about whether the tone was a standard or deviant. Negative co-I values indicate synergy: information about stimulus identity is available from the joint pattern across two time points that is not present in either time point alone. This measure was estimated with the same Gaussian-copula approach used for MI ^32^.

Co-I was computed separately for each channel and condition, and the condition co-I difference was defined as Unattended-minus-Attended. The co-I summary is shown in Fig. 4, with condition-specific co-I matrices in panels **a,b,e,f**, condition differences in panels **c,g**, and participant-level statistical significance summaries in panels **d,h**.

**Figure 4:**
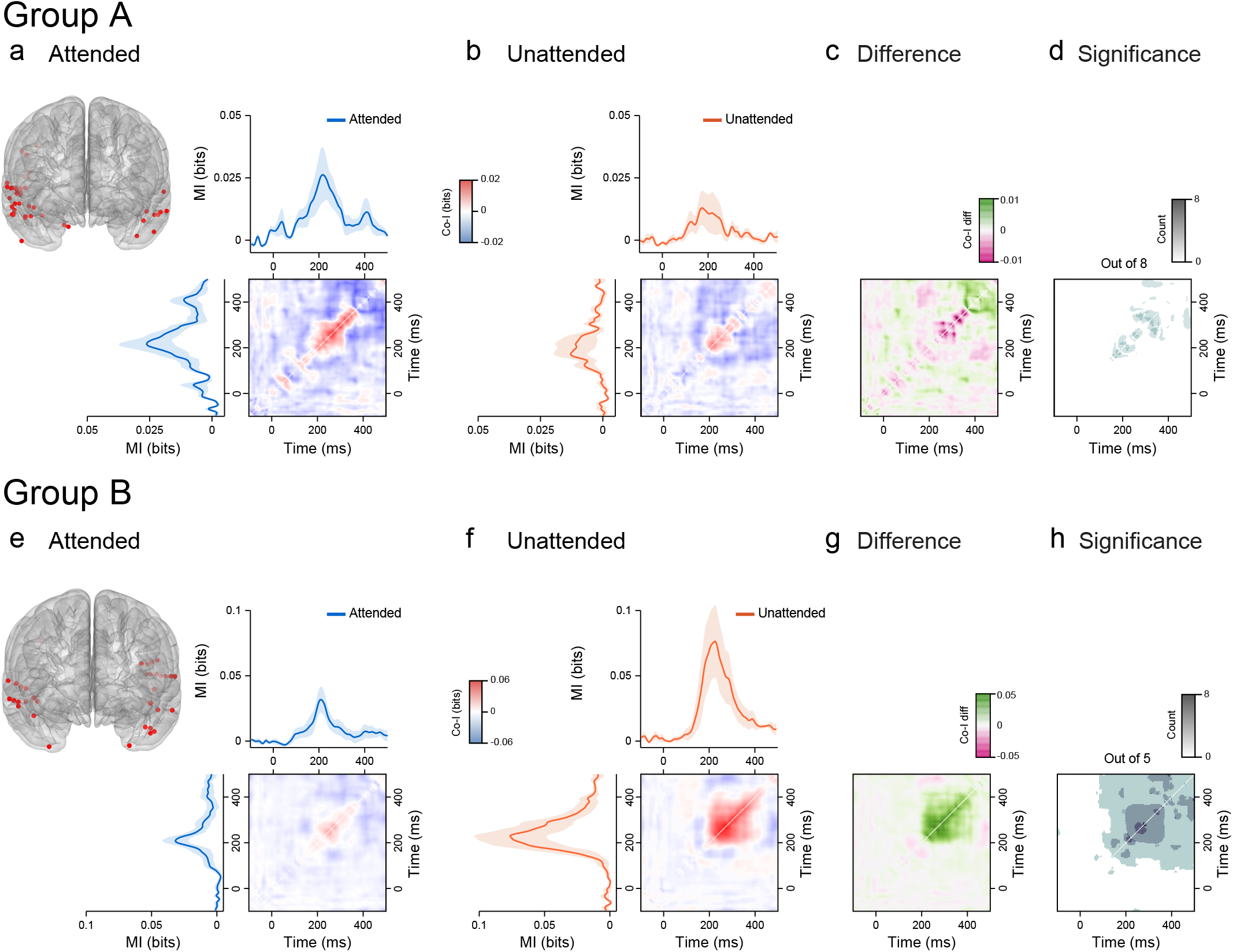
Empirical temporal co-information in Group A and Group B. Co-information (CoI) was computed between pairs of response time points for temporal electrodes with significant attention-related MI modulation. Positive CoI (plotted in red) indicates redundant temporal coding, meaning that overlapping prediction-error information is carried at two response time points. Negative CoI (plotted in blue) indicates synergistic temporal coding, meaning that prediction-error information is available from the joint response across two time points more than from either time point alone. Panels **a–d** show Group A and panels **e–h** show Group B. Attended and Unattended panels (**a,b,e,f**) show temporal electrode locations, condition-specific MI traces, and condition-specific CoI matrices. Difference panels (**c,g**) show Unattended-minus-Attended CoI. In these difference maps, green indicates a positive shift in CoI, corresponding to relatively more redundant or less synergistic temporal coding in the Unattended as compared to the Attended condition, whereas magenta indicates a negative shift in CoI, corresponding to relatively more synergistic or less redundant temporal coding in the Unattended compared to the Attended condition. Significance panels (**d,h**) show, for each time-by-time coordinate, the number of participants with significant condition-related CoI modulation after participant-level stimulus-label shuffling and max-statistic correction over the CoI matrix (out of 8 or 5 participants). Scale bars show the displayed ranges for condition CoI, CoI difference, and participant-count significance maps.

At a broad level, the empirical data showed a structured co-I pattern, with redundancy concentrated around response peaks and synergy linking earlier and later time points, consistent with previous findings of PE information dynamics ^14,15^. Empirical Group A showed a broad shift of both redundant and synergistic regions toward zero between the Attended and Unattended conditions: synergistic time points became less synergistic and redundant time points became less redundant, consistent with an overall reduction in temporally distributed PE information under distraction (Fig. 4**c,d**).

Empirical Group B showed a qualitatively different pattern. While synergistic time points also became less synergistic (change toward zero), redundant time points became *more* redundant (change away from zero; Fig. 4**g,h**). This can be seen in the co-I difference contours: whereas Group A shifts redundant time points toward zero (less redundancy; Fig. 4**c**), Group B shifts them away from zero (more redundancy; Fig. 4**g**). This qualitative distinction was not apparent in the synergistic time points, which changed in a similar direction to Group A (toward zero). In short, empirical Group B showed a shift in the balance of temporal co-I structure, with reduced synergy but stronger redundancy in the Unattended condition.

These results indicate that the transition to the Unattended condition shifted the balance of temporally distributed prediction-error information differently across groups. In Group A, the information balance moved toward zero, consistent with a weakening of distributed prediction-error information. In Group B, the information balance shifted toward reduced synergy but relatively stronger redundancy in the Unattended condition, counterintuitively indicating stronger redundant encoding of PE-information during the Unattended part of the experiment

### Temporal MI dynamics explain the empirical MI and co-I regimes

The empirical MI and co-I analyses established that Attended and Unattended conditions differed not only in the magnitude of PE information, but also in how the content of that information was organised across different time points within the trial.

The next question was then why these differences between conditions appeared. One possibility is that attention produced a stable shift between tasks. Another is that the apparent difference between conditions reflected the history of learning across the experiment, because the roving oddball sequence allows expectations to build and change over repeated stimulus exposure ^36–38^. This distinction was especially important because the Unattended condition always followed the Attended condition.

To characterize temporal dynamics, we computed MI in partially overlapping windows of 30 trials (50% overlap; 15-trial step) spanning the full experiment, from the start of the Attended condition to the end of the Unattended condition. Within each window, we extracted the peak MI, defined as the MI value obtained at the time point of maximum MI, yielding one value per window and participant.

We then fitted linear mixed-effects models to these peak MI values, with fixed effects for Group, Window, and their interaction, and random effects for Subject. We compared three candidate models. The condition mean-shift model tested whether there was a mean difference in MI between the Attended and Unattended halves of the experiment, without modelling a progressive change within either half. The linear model tested whether MI changed gradually across the full experiment. Finally, the hinge model tested whether MI followed a single trajectory during the Attended portion and then changed trajectories at the Attended-to-Unattended transition (window τ = 7; see Methods).

In the thalamus, the condition mean-shift model provided the best overall account of the data by AIC (condition mean-shift: AIC = -317.04; linear: AIC = -311.47; hinge: AIC = -309.98), indicating lower MI in the Unattended than in the Attended half of the experiment (β = −0.0368, *P* = 0.0069; Fig. 5**a**). Thus, thalamic MI was better characterised as a stable shift between task halves than as a gradual change over time, or as a shift in slope trajectories at the attentional manipulation.

**Figure 5:**
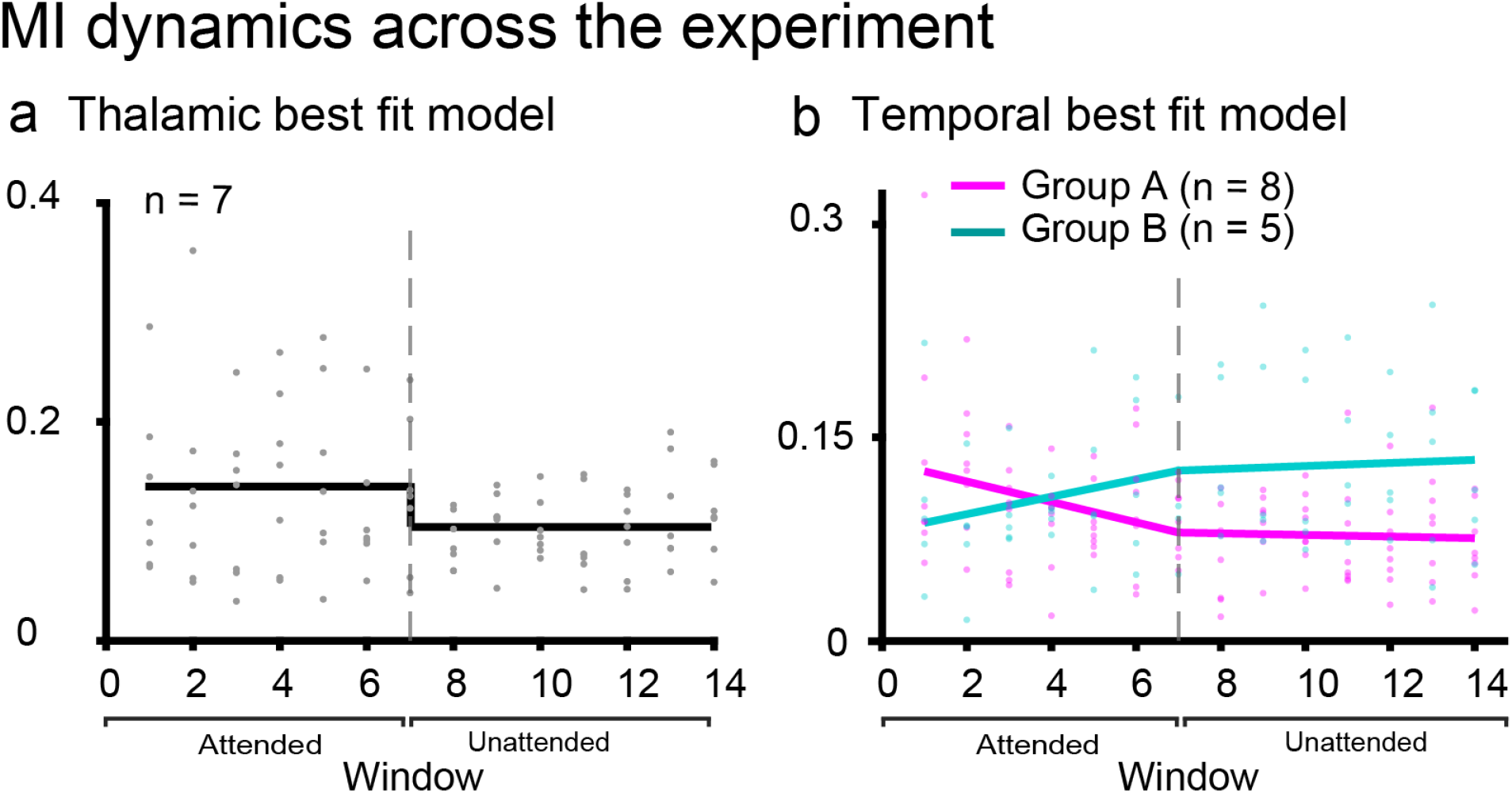
Peak MI across the experiment in thalamic and temporal electrodes. Peak mutual information (MI; bits) was computed in overlapping 30-trial windows (15-trial step) and plotted against window index. Each dot shows an individual subject’s peak MI for a given window. Solid lines show fixed-effect predictions from the selected linear mixed-effects model for each group (Attended, blue; Unattended, orange; sample sizes indicated in the legend). Brackets beneath the x-axis denote the Attended and Unattended phases. For thalamic electrodes (left), AIC selected a condition mean-shift model, capturing a stable difference between task halves. For temporal electrodes (right), the hinge model shown here was supported by LRT, residual bootstrap, and AIC; the dashed vertical line marks the prespecified transition between task halves.

Temporal-cortex MI showed a different pattern (Fig. 5**b**). Here, models that allowed MI to evolve over time provided a better account of the data, and the hinge model was supported over the linear model by the likelihood-ratio test (LRT), residual-bootstrap test, and AIC (hinge: AIC = -703.67; linear: AIC = -699.63; LRT: χ^2^(2) = 8.04, *P* = 0.018; bootstrap *P* = 0.015). This pattern is not consistent with a simple increase or decrease in attention in MI, nor with a single monotonic drift across the experiment. Instead, temporal MI changed trajectory around the Attended-to-Unattended transition.

Before the transition, Group A showed decreasing MI whereas Group B showed increasing MI (Group × Window: β = 0.01334, s.e. = 0.00300, *t* = 4.44, *P* < 0.001; Fig. 5**b**). After the transition, both trajectories flattened, with no reliable slope in either group.

These results indicate that the temporal MI difference between groups was not simply a static difference between Attended and Unattended task states. Instead, the two temporal-cortex groups followed opposite learning-related trajectories during attended listening, consistent with the expectation-building dynamics of roving oddball stimulation ^36–38^, and these trajectories were interrupted or stabilised after the transition to the visual task (Fig. 5**b**). Thus, temporal PE information appears to reflect a dynamic learning regime that is sensitive to attentional state, rather than a uniform increase or decrease in attentional gain.

Keeping the original contrast-defined Group A/B labels fixed, we next tested whether the same groups differed when each condition was analysed in isolation. During the Attended half, Group A showed a negative mean temporal MI slope whereas Group B showed a positive mean slope, with an observed Group B-minus-Group A slope difference of 0.01047 and a two-sided exact permutation result of *P* = 0.047 across all 1287 same-sized group relabellings. By contrast, during the Unattended half, the corresponding slope difference was small and not reliable (Group B-minus-Group A slope difference = -0.00218; exact permutation *P* = 0.450). Thus, the original groups differed specifically in their Attended temporal MI trajectories, whereas their Unattended trajectories were statistically indistinguishable.

As a confirmatory analysis, we repeated the temporal-cortex analysis using a grouping defined by the sign of each participant’s attended pre-transition temporal MI slope (windows 1– 7), with participants with negative slopes assigned to confirmatory Group A and those with positive slopes to confirmatory Group B. This attended-slope grouping closely recapitulated the original participant partition (12/13 participants matched). The key result was preserved and strengthened: for windowed peak MI, the hinge model again outperformed the linear alternative by LRT, residual bootstrap, and AIC (hinge versus linear: ΔAIC = -8.72; LRT: χ^2^(2) = 12.72, *P* = 0.0017; bootstrap *P* < 0.001), indicating that confirmatory Groups A and B showed opposing trajectories during the Attended portion of the experiment that flattened after the Attended-to-Unattended transition.

Further, to test that the hinge-like cortical MI effect did not depend on arbitrary analytical choices (i.e., window size/overlap), we repeated the temporal analysis across window sizes of 20, 30, and 40, each evaluated at 25%, 50%, and 75% overlap (a 3 × 3 grid; 9 window settings total). Across the two grouping schemes (Group A/B and positive/negative slope), the linear-versus-hinge LRT was significant in 15/18 analyses (Group A/B: 6/9; positive/negative slope: 9/9), and the residual-bootstrap test was significant in 14/18 analyses (Group A/B: 6/9; positive/negative slope: 8/9). By AIC, the hinge model was preferred in 13/18 analyses. These results indicate that the temporal hinge-like regime change remained robust across a balanced range of window and overlap choices.

As an additional confirmatory, data-driven analysis, participant-normalized full temporal MI trajectories were entered into an unsupervised two-cluster solution. The resulting clusters largely replicated both the original peak-based grouping and the attended-slope grouping (8/13 and 9/13 participants matched, respectively, after label alignment), and again strongly favoured a hinge over a linear model by LRT, residual bootstrap, and AIC (hinge versus linear: ΔAIC = -13.33; χ^2^(2) = 17.33, *P* = 1.7 × 10^−4^; bootstrap *P* < 0.001).

Together, these convergent analyses show that the hinge-like temporal-cortex regime change reflects a strong and reproducible dynamic in the cortical MI data. The effect was recovered under the attended-slope grouping, remained robust across a balanced grid of window sizes and overlap, and was also recovered by the trajectory-clustering analysis. This indicates that the hinge captures a stable organizational feature of the data across complementary choices of grouping and windowing.

In summary, thalamic MI was best explained by a simple condition mean shift by AIC, with lower MI in the Unattended than in the Attended half of the experiment (Fig. 5**a**). Temporal-cortex MI, by contrast, was not well described by a simple condition mean shift, but instead exhibited distinct dynamics across both condition and group: Group A and Group B showed opposite trajectories during the Attended portion of the experiment, and both trajectories flattened after the Attended-to-Unattended transition (Fig. 5**b**). In the temporal analysis, this hinge-like profile was supported by LRT, residual bootstrap, and AIC across a wide variety of confirmatory analyses. Together, these results indicate that thalamic and temporal-cortex MI dissociate across the experiment: thalamic MI reflects a stable shift in attentional state, whereas cortical MI captures a robust condition-by-group regime change in the dynamics of PE representations.

### A brain-constrained model explains temporal MI and co-I regimes

The temporal-cortex MI trajectories suggested that the Group A/B dissociation was not simply a static difference in attentional gain. Instead, MI evolved over repeated roving-oddball exposure: during attended listening, Group A and Group B followed opposing trajectories, whereas both trajectories flattened once attention was diverted (Fig. 5**b**). This temporal structure points to a learning-dependent effect. In a roving oddball sequence, each deviant becomes the next standard through repetition, so changes across windows are naturally interpreted as changes in the formation and updating of auditory expectations rather than as differences in physical stimulus drive alone. This interpretation is consistent with predictive-processing accounts in which prediction errors update sensory expectations over time ^1–4^, and with evidence that repeated standards strengthen auditory memory traces and shape subsequent mismatch responses ^36–38^. It also aligns with recent intracranial evidence showing that oddball representations of standard on deviant stimuli in the hippocampus can progressively separate over minutes, demonstrating that oddball learning can be expressed as evolving neural discriminability over time in intracranial recordings, even during unconsciousness ^31^. In the present data, however, this learning-like temporal evolution was observed in the temporal neocortex and was interrupted when attention was diverted, suggesting an attention-sensitive cortical learning process.

In particular, the attended phase may have allowed repeated standards and deviants to drive Hebbian updating of stimulus-specific expectations differently in Groups A and B. This idea is consistent with predictive-processing accounts in which prediction errors update sensory expectations over time ^1–4^, because repeated exposure to standards and deviants provides the conditions for such expectations to build and change across the experiment ^36–38^.

Thus, we hypothesised that the differences in temporal-cortex MI trajectories between Groups A and B might be caused by different levels of attention or attentional capacity between the groups. Specifically, because Group A encoded more PE information at the beginning of the experiment, we hypothesised that they exhibited higher levels of attention. In contrast, because Group B encoded less PE information at the beginning of the experiment, we hypothesised that they exhibited lower levels of attention.

We therefore used a brain-constrained neurocomputational model to test the hypothesis that different effective attentional regimes can alter learning-dependent response trajectories while holding the sensory input and network architecture fixed. Specifically, we asked whether attention-related changes in circuit state could modulate the Hebbian learning induced by attended roving-oddball exposure, thereby producing divergent temporal trajectories in ERP, MI, and co-I signatures that resemble the empirical Group A/B difference. If this were the case, it would identify attention-modulated Hebbian learning as a potential mechanism to explain the differing MI trajectories of empirical groups A and B.

To this end, we applied a brain-constrained neurocomputational model of fronto-temporal perisylvian cortices ^39^ that has previously been used to explain the neural mechanisms underlying attention-dependent cortical responses, automatic auditory change detection and ERP responses, endogenous decision dynamics, verbal working memory, and prediction-error information dynamics ^14,40–42^. The network comprises six reciprocally connected cortical areas: three auditory-temporal areas (A1, AB, PB) and three frontal areas (PF, PM, M1). Critically for the present question, the model implements ongoing Hebbian synaptic plasticity during stimulus exposure, making it suitable for testing whether attention-like changes in circuit state can reshape the evolution of learning rather than merely rescale evoked responses. Within this architecture, attention-like control can be approximated by changing both global inhibitory regulation and the strength of between-area connectivity. We therefore operationalised the two model regimes by varying global inhibition (GI) and between-area connectivity (Ffb), while leaving the sensory input, architecture, and all other parameters unchanged (see Methods).

We ran 8 independently initialised simulations for Group A and 5 independently initialised simulations for Group B, matching the empirical group sizes. In model Group A, lower global inhibition (GI = 90) and stronger between-area connectivity (Ffb = 750) implemented a higher-attention regime. In model Group B, higher global inhibition (GI = 120) and weaker between-area connectivity (Ffb = 500) implemented a lower-attention regime (Fig. 6**a**). Each simulation comprised 240 trials, matching the empirical experiment. These simulations did not include an Attended-to-Unattended switch midway through the run; rather, they instantiated two fixed-parameter regimes designed to test whether stable differences in attentional state during tone listening could reproduce the group-specific empirical dynamics observed in the Attended condition. We recorded simulated firing-rate responses from the three temporal layers (A1, AB, PB), derived proxy ERP responses from layer-averaged activity, and then applied the same information-theoretic analyses used for the empirical data.

**Figure 6:**
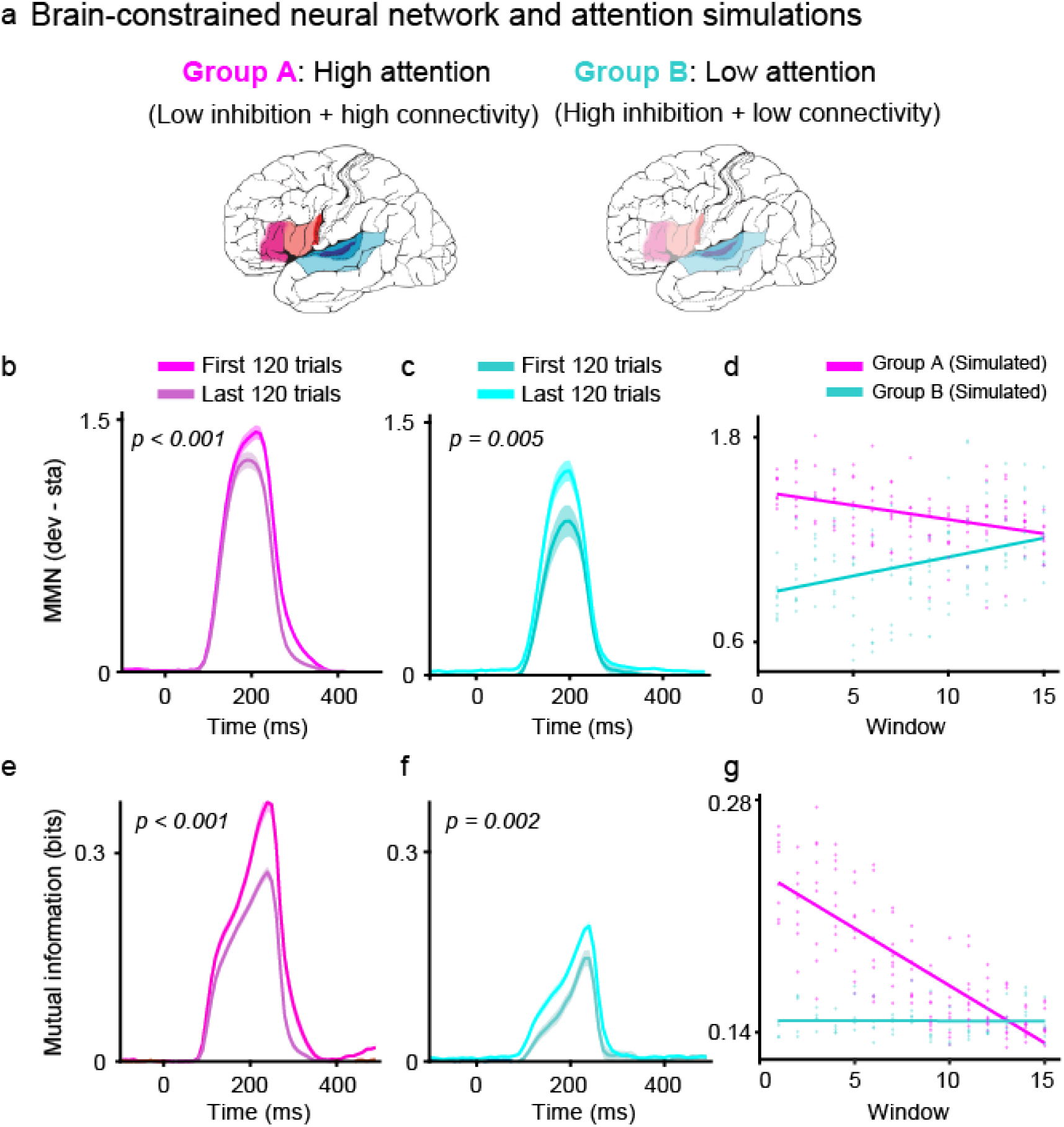
Model mutual information across the experiment. (a) Brain-constrained fronto-temporal model and attention-regime manipulation. Group A was simulated as a higher-attention regime with lower global inhibition (GI = 90) and stronger between-area connectivity (Ffb = 750), whereas Group B was simulated as a lower-attention regime with higher global inhibition (GI = 120) and weaker between-area connectivity (Ffb = 500). (b–c) Model event-related potential difference (ERP; DEV – STD) for the first 120 and last 120 trials in Group A and Group B. (d) Windowed ERP trajectories across the 240-trial simulation. (e–f) Model mutual information (MI; bits) for the first 120 and last 120 trials in Group A and Group B. (g) Windowed MI trajectories across the 240-trial simulation. Group A used 8 independently initialised simulations and Group B used 5 independently initialised simulations, matching the empirical group sizes. Shaded bands indicate ± SEM across simulations; time is shown relative to stimulus onset (0 ms).

The model reproduced the main group-specific first-versus-last patterns in MI and ERP (Fig. 6**b,c,e,f**). Across these simulations (Group A: *n* = 8; Group B: *n* = 5), simulated Group A showed higher temporal peak ERP in the first 120 trials than in the last 120 (paired *t*(7) = 6.18, *P* < 0.001, *d*_*z*_ = 2.18, 95% CI [0.209, 0.469]), and temporal peak MI showed the same direction of effect (*t*(7) = 5.73, *P* < 0.001, *d*_*z*_ = 2.03, 95% CI [0.061, 0.147]). Simulated Group B showed the opposite ERP pattern, with higher temporal peak ERP in the last 120 trials than in the first (*t*(4) = −5.46, *P* = 0.0055, *d*_*z*_ = −2.44, 95% CI [-0.428, -0.139]). Temporal peak MI in Group B showed the same direction of effect (*t*(4) = −7.61, *P* = 0.0016, *d*_*z*_ = −3.40, 95% CI [-0.064, -0.030]).

The locus of modulation differed by layer: simulated Group A showed the strongest MI modulation in higher temporal layers (AB, PB), with minimal modulation in A1, whereas simulated Group B showed modulation primarily in A1, with little reliable change in AB/PB (Supplementary Fig. S1).

We next asked whether the model also reproduced the empirical temporal-cortex trajectories across the full simulation. We computed windowed MI and ERP across the simulation time course (window size 30, step 15) and fitted linear mixed-effects models. For MI, the simulated data captured the Group A pattern well: cortical peak MI declined significantly across windows (Window: β = − 0.0145, *P* < 0.001; Fig. 6**g**). Although the Group B trajectory differed significantly from Group A (Group × Window: β = 0.0139, *P* < 0.001), the simulated Group B slope was approximately zero, indicating a largely flat trajectory rather than the positive increase observed empirically. Thus, the model reproduced the temporal-cortex MI dynamics of empirical Group A, but not those of empirical Group B.

The simulated ERP trajectories reproduced the empirical group divergence more completely than the simulated MI did. Peak ERP declined significantly across windows in Group A (Window: β = −0.0176, *P* < 0.001; Fig. 6**d**), whereas Group B showed a significantly more positive slope than Group A (Group × Window: β = 0.0451, *P* < 0.001), corresponding to an increasing ERP trajectory rather than the decline observed in Group A. Thus, whereas simulated MI captured the empirical Group A trajectory but not the increasing Group B trajectory, simulated ERP reproduced the expected between-group divergence across both groups.

Finally, we asked whether the brain-constrained model used to simulate different attentional regimes could also capture the qualitative changes observed in the empirical results; does the model co-I evolve similarly to the empirical co-I?

To this end, we applied the same co-I analysis to the model data as to the empirical data, and then compared the resulting model co-I-difference maps with the empirical maps (Fig. 7). For the empirical data, the contrast was Unattended-minus-Attended; for the simulated data, the contrast was last-120-minus-first-120 trials. The co-I difference matrices were compared using two complementary metrics: structural similarity (SSIM; previously used in Gelens et al. ^14^), which quantifies overlap in the layout of significant co-I modulation within the 0–500 ms comparison window, and signed Pearson correlation, which quantifies whether corresponding regions changed in the same or opposite directions.

**Figure 7:**
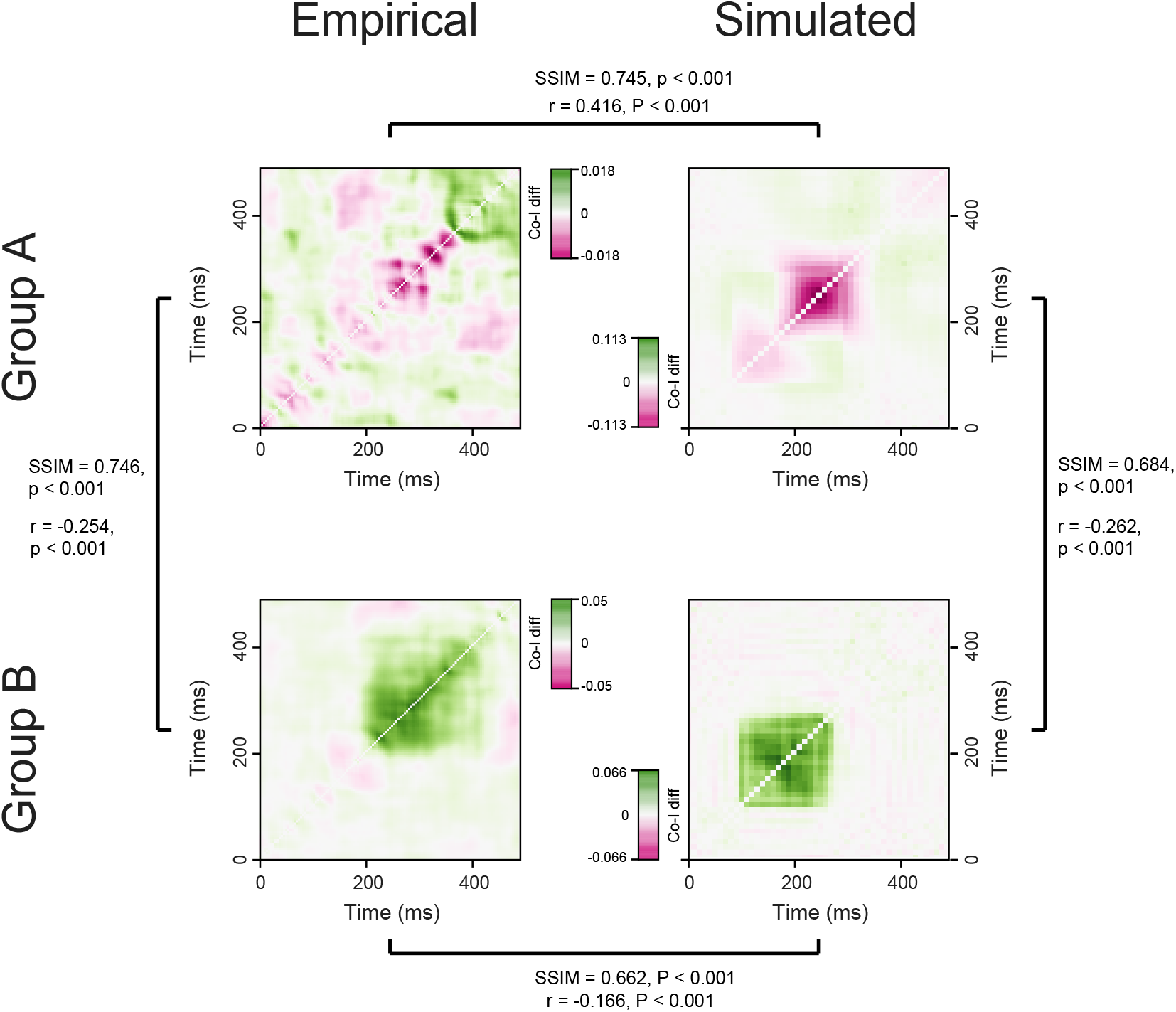
Empirical and simulated co-I difference matrices. The empirical panels show the same type of Unattended-minus-Attended co-i difference maps as in the empirical co-I figure (Fig. 4**c,g**), replotted here for direct comparison with the model. The simulated panels were generated with the same CoI analysis pipeline, but using the contrast between the last 120 and first 120 simulated trials. Columns show empirical and simulated data; rows show Group A and Group B. Green indicates a positive CoI change, corresponding to a shift toward more redundant temporal coding, whereas magenta indicates a negative CoI change, corresponding to a shift toward more synergistic temporal coding. For visualization, each matrix is plotted with its own colour limits to show its internal structure clearly. For the SSIM analyses matrices were rendered with shared colour limits before comparison. Brackets indicate the matrix pairs compared statistically: empirical versus simulated within each group, and Group A versus Group B within empirical or simulated data. Each bracket reports structural similarity (SSIM), quantifying similarity in the spatial layout of CoI modulation, and signed Pearson correlation (*r*), quantifying whether corresponding matrix regions changed in the same or opposite direction.

We first used these metrics to compare the empirical Group A and Group B co-I-difference maps to test whether the two empirical regimes shared a common spatiotemporal modulation layout while differing in signed direction. We then compared each empirical group with its corresponding simulated group to test whether the model reproduced the empirical co-I

modulation pattern within each regime. Finally, we compared simulated Group A with simulated Group B to test whether the model itself generated the same between-regime divergence observed in the empirical data. Significance was assessed against shuffle-based null distributions with Bonferroni correction across the four planned comparisons.

Within the empirical data, Group A and Group B did not simply differ in the magnitude of co-I modulation; rather, they differed in how the balance of redundant and synergistic information in the brain was modulated (Fig. 7**a,c**). The two groups showed significant structural similarity (SSIM = 0.746, Bonferroni-corrected *P* < 0.001), indicating that they shared a non-random scaffold of modulation across the co-I matrix. Crucially, however, the signed Pearson correlation between these matrices was negative (*r* = −0.254, Bonferroni-corrected *P* < 0.001), showing that corresponding regions of this shared scaffold tended to shift in opposite signed directions across groups. Thus, the empirical difference was not merely quantitative, that is, not just a stronger versus weaker version of the same effect, but qualitative: the same underlying co-I architecture was reorganized in opposite ways in Group A and Group B.

The model partly reproduced these empirical co-I regimes. Simulated Group A closely matched empirical Group A both in structural organisation (SSIM = 0.745, Bonferroni-corrected *P* < 0.001) and in signed directionality (*r* = 0.416, Bonferroni-corrected *P* < 0.001; Fig. 7**a,b**), indicating that, as with the empirical data, the model captured not only where co-I modulation occurred, but also how the balance of redundant and synergistic information shifted differently between the two groups.

Empirical Group B and simulated Group B also showed significant structural similarity (SSIM = 0.662, Bonferroni-corrected *P* < 0.001), but their signed correlation was negative (*r* = −0.166, Bonferroni-corrected *P* < 0.001, opposite-direction test; Fig. 7**c,d**). Thus, the model reproduced part of the spatial layout of Group B co-I modulation, but not its dominant signed pattern.

Finally, simulated Group A and simulated Group B were themselves highly similar in structure (SSIM = 0.684, Bonferroni-corrected *P* < 0.001) but opposite in signed directionality (*r* = −0.262, Bonferroni-corrected *P* < 0.001; Fig. 7**b,d**), mirroring the divergence observed in the empirical groups.

Taken together, these findings suggest that the critical group difference lies not in a common co-I pattern, but in how that architecture is driven across conditions. Group A and Group B appear to engage a similar spatiotemporal pattern of co-I modulation, yet express it in opposite directions. The model successfully captured this logic for Group A, but only partially for Group B, possibly due to a temporal lag between the empirical co-I and the model (see Fig. 7). These results support the idea that the empirical groups are distinguished by qualitatively different informational regimes rather than by a simple difference in effect strength, which are replicated by the model.

## Discussion

In the present study, we examined how PE information encoded in intracranial ERPs from thalamic and temporal regions is modulated by attention. Our main finding is that attentional modulation of PE information was not uniform across the auditory hierarchy. Thalamic PE information was best characterised by a relatively sustained state-dependent shift between task contexts. By contrast, temporal-cortex PE information showed pronounced inter-individual heterogeneity: one subgroup showed reduced PE information after attention was diverted, whereas another showed the opposite pattern. Crucially, this temporal heterogeneity was not simply a static attentional gain effect. Instead, Group A showed a decrease in temporal MI during attended listening, Group B showed an increase during the same period, and both trajectories flattened after the transition to the visual (the unattended condition) task (Fig. 5**b**). Thus, the critical temporal effect was a change in the evolution of PE information during repetition-dependent learning in the attended condition.

This pattern helps reconcile inconsistent findings in the literature on attentional modulation of mismatch responses. Previous studies have reported enhancement of prediction-error or ERP-like responses under attention, attenuation under competing visual load, or little group-level change despite substantial individual variability ^19,27–29^. Our findings fit predictive-processing accounts in which attention and learning precision-weight PE signals, i.e., change their effective gain and impact on hierarchical updating rather than simply scaling mismatch amplitude ^2,16,17,19,43^. Within this framework, as the brain-realistic model’s simulation results suggest, one reason for the apparent heterogeneity in previous work may be that attention interacts with ongoing repetition-dependent learning, such that the direction and magnitude of the effect depend on the listener’s attentional capacities and learning regime (see below).

### Thalamic PE information reflects sustained state-dependent modulation

Various empirical findings have implicated thalamic circuits in auditory deviance detection and prediction-error processing, indicating that mismatch-related signals are not generated exclusively in cortex ^5,44,45^. Related human single-neuron evidence has also reported auditory deviance responses outside the thalamus, including in the amygdala, hippocampus, insula, and auditory cortex, further emphasizing that mismatch-related responses can be distributed across broader temporal-limbic and cortical circuits ^46^. In the present data, thalamic PE information showed a pattern distinct from the temporal cortex. Although significant thalamic modulation was observed only in a subset of patients (6/12), its direction was more consistent across individuals: most patients with significant thalamic effects showed higher MI in the Attended than in the Unattended condition. In addition, unlike temporal-cortex MI, thalamic MI did not exhibit a hinge-like change in slope at the task switch, but was better characterized by a sustained shift between task contexts. Thus, the heterogeneity that dominated temporal PE information was not mirrored in the thalamus.

One interpretation is that thalamic MI indexed a broader change in thalamocortical state between the two task contexts, for example, in gain regulation, routing, or attention-linked control, rather than a dynamically evolving learning process. This interpretation is consistent with prior work suggesting that thalamic attentional effects often reflect broader state-dependent control of cortical processing, including gain regulation, gating, and arousal-linked modulation. ^47–50^. It is also consistent with work showing that auditory thalamic neurons exhibit robust stimulus-specific adaptation, and that this history-dependent sensitivity is shaped locally within thalamic circuitry rather than simply inherited from cortex ^51–54^. At the same time, the anatomical interpretation of this effect should be approached with caution. Much of the strongest direct evidence for thalamic deviance processing comes from auditory thalamic nuclei such as the medial geniculate body and related non-lemniscal auditory thalamic circuits ^5,44,45^, whereas the significant thalamic contacts in the present dataset were located outside classical auditory thalamus. Thus, our effect is best interpreted as reflecting a broader thalamic contribution to state-dependent attentional gating rather than a nucleus-specific auditory-thalamic mechanism. However, as a caveat, the deviant-minus-standard contrast in a roving oddball task is unlikely to isolate prediction error from stimulus-specific adaptation. Thus, the thalamic effect could also partly reflect attentional modulation of stimulus-specific adaptation.

### Temporal-cortex PE information reflects distinct learning regimes during attended listening

Unlike thalamic MI, the grand average across all patients showed that temporal MI did not differ substantially between the Attended and Unattended parts of the experiment. However, this absence of an overall mean effect masked pronounced interindividual heterogeneity. Among patients showing temporal modulation, one subgroup showed decreased PE-information encoding in the Unattended relative to the Attended condition, whereas another subgroup showed the opposite pattern. This kind of heterogeneity is easy to miss in grand averages and is precisely the kind of effect for which subject-level prevalence approaches are informative ^33^. It is also consistent with recent ERP work showing that even when group-level attentional effects are small or absent, individual participants can exhibit both attention enhancement and suppression ^28^. Thus, what might appear as a weak or absent mean effect at the group level can conceal robust but opposing subject-level regimes.

Crucially, the temporal subgroup difference was not simply due to the attentional manipulation shifting PE information up or down between conditions. Rather, the two groups were distinguished by different MI trajectories already during the Attended part of the experiment: before the Attended-to-Unattended transition, Group A showed decreasing temporal MI across windows, whereas Group B showed increasing temporal MI. After the transition to the visual task, both trajectories flattened. Thus, the primary divergence occurred during attended listening, and this shift in trajectory was attenuated when attention was diverted. This hinge-like pattern was recovered with a confirmatory regrouping based on attended pretransition MI slope and by a fully data-driven clustering analysis of temporal MI trajectories, and remained robust across a balanced sweep of window sizes and overlaps.

In the roving oddball paradigm, each new tone first acts as a deviant and then becomes the new standard through repetition. Changes across trials, therefore, primarily reflect the formation and updating of auditory expectations rather than differences in physical stimuli ^36^. Repeated standards are known to strengthen the sensory memory trace and to shape subsequent mismatch responses ^37,38^. As model’s predictions emerging from much earlier simulation results had originally suggested ^40^, because the temporal cortex is involved in auditory regularity learning and deviance detection, temporal-cortex MI may index the evolving comparison between incoming tones and learned auditory expectations, rather than only low-level repetition adaptation ^55^. Accordingly, the Group A/B difference is best interpreted as reflecting distinct learning and/or attentional regimes during the attentive listening condition. Attention, therefore, appeared to shape how PE information evolved across attended repetition, rather than simply shifting PE information up or down between conditions.

These findings are consistent with recent results demonstrating PE learning in the human hippocampus ^31^. However, our findings reveal a distinct learning regime in the temporal cortex. While hippocampal learning has been shown to occur in unconscious patients ^31^, our results indicated that learning in the temporal neocortex was attention-dependent and disrupted when attention was diverted from the oddball stimuli. Interestingly, we also found that thalamic PE-encoding showed no evidence of learning throughout the experiment. Together, our findings indicate that PE-learning across the predictive hierarchy is not governed by a single relationship between stimuli and learning, but rather depends on consciousness and attention in ways that vary with the neuroanatomical circuit and area.

However, as the cohort consisted of patients undergoing invasive monitoring for drug-resistant epilepsy, these individual differences should also be interpreted with clinical heterogeneity in mind. Statistical learning engages medial-temporal and hippocampal circuitry, including during implicit auditory structure learning, and statistical-learning performance in epilepsy has been linked to seizure frequency and hippocampal subfield volume ^56,57^. In addition, antiepileptic medication can affect attention, processing speed, and memory ^58^. Factors such as seizure focus, medial-temporal or hippocampal pathology, and medication could therefore influence statistical learning and may contribute to the distinct temporal regimes of the groups.

Moreover, in roving oddball paradigms, ERP mismatch responses and the deviant-standard contrasts used to quantify them are not process-pure: they can reflect prediction-error-related updating together with stimulus-specific adaptation and other history-dependent differences between standards and deviants ^5,55^ Thus, the differences between the groups might also partially reflect differences in stimulus-specific adaptation.

### A brain-constrained model mechanistically explains the observed data

We hypothesized that the two distinct temporal learning trajectories could reflect differences in effective attention during the experiment’s attended phase, or, more generally, differences in attentional capacity across patients. This interpretation is supported by the hinge-model result, which showed that both subgroup trajectories were flattened when attentional demands increased during the visual task, suggesting that the dynamics ongoing in both groups require attentional resources. To test this idea, we used a brain-constrained model to simulate the task’s attended condition, keeping the sensory input and all non-attention parameters constant, and manipulated global inhibition and between-area connectivity strength as proxies for attention in the model. This manipulation extends the original Garagnani et al.’s model implementation in which the global-inhibition (GI) parameter alone was the independent variable (low GI modeled a high-attention regime and higher GI a low attentional-resources condition in simulated word and pseudoword responses – see Supplementary Fig. S2; ^39^) by pairing GI with between-area connectivity changes in the current simulations. Crucially, the current work goes well beyond all previous modeling studies with this neural architecture, as it investigates the changes induced by learning in the (simulated) cortical responses over a given time span. This was done by comparing ‘early’ vs. ‘late’ network responses to novel (simulated) auditory stimuli (previous works only compared responses to well-known, learnt items – taken as model correlates of familiar words – to randomly scrambled and recombined versions of such stimuli, representing unknown pseudoword items ^39,40^), thus investigating the effects of learning under different attentional regimes on the responses to initially new sounds as they become more “familiar” to the network.

The neurocomputational model we used has been widely employed to study and explain neurophysiological responses to familiar and unfamiliar auditory stimuli. Earlier work with this architecture showed that attention-like changes in global inhibition can qualitatively alter ERP-like responses ^39^, that auditory change-detection effects can only be fully explained when Hebbian learning mechanisms (leading to the emergence of memory circuits) are added to adaptation and inhibition mechanisms ^40^, and that the same anatomy-and-plasticity principles naturally generate persistent activity – the hallmark of working memory – in higher temporal and prefrontal association areas after learning ^59^. More recently, Gelens et al. ^14^ applied this brain-constrained neural architecture to model PE processing and showed that it reproduces and explains synergistic PE coding as a result of recurrent and long-range feed-back/feedforward interactions.

Our simulations produced ERP dynamics qualitatively similar to both empirical groups and reproduced the MI dynamics for Group A, and partially for Group B (see Fig. 6**b–g**). Specifically, in the present data, the key phenomenon to explain was not simply a difference in the overall amount of PE information, but a difference in its temporal evolution during attentive listening. In fact, throughout the Attend condition, Group A showed decreasing temporal MI across windows, whereas Group B showed increasing temporal MI; both trajectories flattened after the Attend-to-Unattend transition. The same qualitative pattern was observed in the ERP dynamics (during the Attend condition), with decreasing PE responses in Group A and increasing PE amplitude in Group B over time. The model captured most of the patterns observed experimentally: it accurately reproduced (and explained – see below) the qualitative difference in ERP changes between Group A and Group B, as well as the significant decrease seen in Group-A’s MI, while the empirical MI increase observed in Group B was not replicated, remaining a “null” effect in the simulation data.

The explanatory account being proposed here for the opposite patterns found experimentally in Groups A and B (decreasing ERP and MI for the former, increasing for the latter) is grounded in a seminal study which used the same neural architecture ^39^, and which suggests that the two seemingly incongruent patterns observed may be just the result of two different levels of attentional resources available (under which learning took place). In fact, as Supplementary Fig. S2 shows, network responses to model correlates of familiar (i.e., word) and unfamiliar (pseudoword) items exhibited opposite patterns depending on the (simulated) level of attention (GI). Specifically, under high attention (top panel), responses to unfamiliar stimuli were *larger* than to familiar ones; with lower attention (bottom panel), the pattern reversed (larger responses to familiar than unfamiliar “sounds”). Importantly, in the present work, where synaptic plasticity mechanisms were constantly and uniformly at work, the ‘familiarity’ of the input stimuli (modelling auditory tones) changed during the experiment, going from initially novel and unfamiliar to well-learned and familiar. According to the original simulations, under higher attentional resources (top panel), the network response to the same stimulus should *decrease* over time (the larger response – in blue – to unfamiliar pseudoword items gradually ‘shrinking’ until it becomes the smaller – red – familiar-word curve). However, the opposite pattern should be observed under lower-attention conditions (bottom panel): as the network learns the repeatedly presented stimulus (i.e., as a corresponding memory trace emerges), the lower blue unfamiliar-item curve gradually ‘grows’ towards the red familiar-item curve. This is exactly what the present simulation results show (Fig. 6**d**), accurately reflecting and explaining the seemingly idiosyncratic pattern observed experimentally (Fig. 5**b**). In short, the model simulations confirmed that the opposite patterns seen experimentally in Groups A and B could be explained by differences in effective attention (or attentional capacity) between the two groups during the experiment’s Attentive-listening task, as hypothesized.

Moreover, and crucially, however, the use of a brain-constrained model, mimicking the structure and neurophysiology of the relevant cortices, also allowed us to formulate a tentative neuromechanistic explanation for the observed experimental data. In particular, the simulations suggest that the cortical mechanisms underlying the interactive effects of attention and stimulus familiarity (directly linked to learning) can be identified in the presence (or absence), and inherent dynamics, of Hebbian cell-assembly circuits, memory traces which putatively emerge in the neocortex as a result of repeated sensory stimulation. As such, associative circuits develop gradually over time; they are absent during the early part of the experiment, but are fully formed by the last trials. Importantly, as explained in the original study ^39^, cell-assembly circuits behave as all-or-nothing functional units that, if sufficiently stimulated, become fully active (“ignite”). Accordingly, the abovementioned larger (or smaller) network responses observed in response to the same (initially unfamiliar, later familiar) input stimuli under different levels of attention can be explained by different interactions among such circuits, as follows. The larger responses – under larger levels of attention – to unfamiliar than to familiar items is a consequence of the fact that an unfamiliar stimulus automatically activates multiple candidate circuits, none of which exactly ‘matches’ the sensory input, but all of which share/respond to *some* of its characteristics. As a result, in the presence of sufficient attentional resources (here, low global inhibition and high between-area connectivity), multiple circuits may simultaneously activate in response to an unfamiliar sensory item, resulting in a response overall larger than that induced by a familiar one (and for which a single, stimulus-specific memory trace exists, is activated, and ignites). Because of the mutually inhibitory dynamics (mediated by the global and local inhibitory mechanisms of the network) that exist between them, however, when stimulated, the multiple memory circuits engage in competitive interactions, whereby they suppress one another (for a detailed explanation, see the discussion in ^39^, Sec. 5.3): if not enough attentional resources are present (i.e., higher GI and lower between-area connectivity), such mutually inhibitory processes prevent *all* of the stimulated circuits from igniting. This results in an overall smaller response than that, again, induced by a familiar item, for which a single, unrivalled memory trace exists and ignites, even when attention is lower.

This explanation is bolstered by additional prior work using the same model architecture. Garagnani and Pulvermüller showed that, within this fronto-temporal network, adaptation and local inhibition can explain some auditory change-detection effects for unfamiliar repeated sounds, but are insufficient to account for the stronger mismatch-like responses elicited by familiar auditory patterns ^40^. In the model, these stronger responses only emerge when Hebbian memory circuits formed through learning are available, indicating that adaptation, inhibition, and Hebbian plasticity make distinct contributions to ERP dynamics. This is particularly relevant to the present findings because the empirical group difference was not a simple shift in the mean PE information between conditions, but rather a difference in how temporal PE information evolved during attended listening, likely signaling learning mechanisms.

Taken together, these findings suggest that the interactive effects of attention and stimulus familiarity (directly linked to learning), modelled here as paired changes in global inhibition and between-area connectivity, are a viable candidate mechanism for the opposing temporal PE-information effects observed across the two patient groups.

The fact that different model regimes altered the temporal evolution of PE information is consistent with the idea that the opposing empirical trajectories may reflect differences in how adaptation, inhibition, and Hebbian plasticity interact during attended listening. Notably,^31^ employed a ‘standard’ recurrent neural network trained with gradient descent (i.e., Backpropagation Through Time, BPTT) to discriminate tone identity and showed that standard and deviant oddball divergence emerged across time steps, replicating their empirical findings in the hippocampus. We employed a more biologically realistic, brain-constrained network here that replicates the anatomical structure and physiological function of the relevant cortices, adding to this body of evidence, expanding on ^31^, and providing a neuromechanistic explanatory account for the observed attention-dependent PE learning within neocortical networks.

### Distraction disrupts the learning-dependent informational architecture of temporal PE representations

The co-I results extend the MI findings by showing that the critical effect was not a simple attentional reshaping of the amount of PE information, but a disruption of the learning-dependent informational signature that formed during attended listening. Earlier work has used co-Information and information decomposition to reveal redundancy and synergy during audiovisual speech perception ^60^, synergistic cross-modal interactions supporting active multisensory decisions ^61^, redundant and synergistic sound coding in mouse auditory cortex ^62^, and dissociable redundant and synergistic interactions during human intracranial reward and punishment learning ^63^. In this context, our co-I results indicate that the temporal signature of PE information was established during attended listening and was subsequently stabilized after the transition to the visual task.

Across both empirical and simulated data, PE responses showed a similar qualitative co-I structure, with redundancy concentrated around the main response peak and synergy linking earlier and later time points, yielding the characteristic off-diagonal early-late synergy observed in earlier studies ^14,15^. This is consistent with findings showing that prediction-error information can be highly synergistic, such that some information about mismatch is available only in the joint relationship between signals or time points ^14^. Importantly, however, the transition to the visual task did not affect this structure uniformly across groups. In Group A, both redundant and synergistic components shifted toward zero, suggesting a broad weakening of the distributed PE representation once the learning dynamics present during attended listening were disrupted. In Group B, synergistic components likewise became less synergistic, but redundant components became more redundant. In the context of the roving oddball paradigm, where repeated standards are known to strengthen the sensory memory trace, this increase in redundancy may be consistent with a more stabilized or persistent standard representation ^37,38,64,65^. Thus, the group difference was not only quantitative (MI: amount of information) but also qualitative (co-I: type of information): perceptual learning during the attended condition altered the balance between redundant and synergistic PE coding in different ways across the two temporal subgroups.

The similarity between empirical and simulated co-I patterns further supports the interpretation that these informational changes are mechanistically meaningful rather than merely descriptive. Notably, while we can manipulate how the model’s ERP responses evolve via paired changes in inhibition and between-area connectivity, we have no control over how the model’s information signature develops over time steps. Thus, the fact that both empirical and simulated co-I signatures evolve in a strikingly similar manner across time steps lends plausibility to the claim that they rely on similar mechanisms. Our results, therefore, suggest that perceptual learning during attended listening helps establish the temporal informational signature of PE representations and that increased visual attentional demands disrupt this process. More broadly, these findings are consistent with the view that separating redundancy from synergy provides a principled way to characterize the informational architecture of brain computations ^14,15,66^.

### Why MI did not capture both simulated trajectories

A likely explanation for the dissociation between simulated ERP and MI, where for Group B ERP-like responses in the model captured the evolving trajectories across time steps but MI did not, is that the two measures are sensitive to different aspects of the response. While ERP reflects an average difference between standard and deviant responses, MI depends on how well these trial-by-trial responses can be separated at the single-trial level ^32^. In the roving simulations, every deviant was preceded by a variable run of 6–10 identical standards (see Methods).

If responses in lower auditory levels of the model are strongly shaped by adaptation to the preceding standard train, then a deviant following 10 standards will elicit a more pronounced mismatch response than a deviant following only 6 standards. Pooling these trials into a single “deviant” category, therefore, increases within-class variability, such that the deviant distribution becomes a mixture of response types with different distances from the standard distribution. Under these conditions, standard and deviant responses can still differ clearly in their averages, thereby producing an ERP effect, while overlapping substantially at the single-trial level, yielding relatively low MI.

This interpretation is particularly plausible for simulated Group B, who showed a dissociation with MI and ERP, and for whom the modulation was concentrated almost exclusively in A1 (model layer 1), and is therefore likely to be more dependent on repetition history and adaptation. By contrast, in simulated Group A, the modulation was expressed more strongly at higher auditory levels, AB/PB, where deviance coding may be less tightly coupled to the exact length of the preceding standard train and more strongly reflects an abstract prediction-error signal. Such layer-specific differences may reduce within-class variability in the deviant responses and make MI more sensitive to the modulation. These factors, combined with the fact that the evolution of the trajectories across the time-steps was computed in 30-trial windows, which yields a noisy MI estimate, might explain why, for group B, MI exhibited a purely flat trajectory.

In this sense, the simulated dissociation between ERP and MI does not indicate the absence of an effect in the Group B regime; rather, it suggests that single-trial informational separability is especially vulnerable to run-length-dependent adaptation in the lower auditory cortex, whereas averaged mismatch responses are not. This interpretation is broadly consistent with both the model architecture, in which adaptation and memory make distinct contributions across cortical levels ^40^, and with empirical work showing that adaptation declines while prediction-error-related processing becomes more prominent higher in the auditory hierarchy ^55^.

## Conclusion

Taken together, these findings suggest that thalamic and temporal PE information follow dissociable but coordinated dynamics. The temporal cortex showed learning-sensitive, highly heterogeneous trajectories, consistent with a role in the formation and updating of auditory regularities during roving stimulation ^36–38,55^. By contrast, thalamic PE information is more consistent across patients and is better captured by a sustained shift between task contexts, in line with broader thalamic contributions to state-dependent gain regulation, attentional gating, and thalamocortical routing ^47–50^.

It is also noteworthy that the reduction in thalamic MI during distraction coincided with the flattening of temporal learning trajectories, suggesting that the thalamic state shift may mark a change in the conditions under which repetition-dependent PE learning can be sustained in the temporal cortex. This temporal correspondence is consistent with predictive-processing accounts in which cortical PE learning unfolds under thalamically mediated changes in gain and control state ^2,16,50^.

Moreover, together with recent evidence for unconscious hippocampal oddball learning ^31^, our findings suggest that PE learning is circuit- and state-dependent: it can persist without consciousness in the hippocampus, but in the temporal neocortex it depends on attention, whereas thalamic PE signals show state-dependent modulation without detectable learning trajectories.

Our findings help explain why studies of attentional effects on ERP have often produced heterogeneous results. Rather than exerting a single, uniform effect on auditory mismatch responses, attention appears to interact with ongoing repetition-dependent learning, so that the same attentional manipulation can attenuate, preserve, or reverse the apparent direction of PE responses depending on the learning regime already present in the listener. This interpretation fits the broader literature, in which some studies report attention-related enhancement of mismatch responses, others report reductions under visual load, and others find little group-level change despite substantial individual variability ^19,27–29^. Our results, therefore, suggest that the long-standing question is not simply whether attention increases or decreases ERP and PE information across different cortical levels, but when and how attention reshapes the learning dynamics that generate PE information in the first place.

## Methods

### Ethics statement

This study was conducted in accordance with the Mayo Clinic Institutional Review Board (IRB 15-006530), which authorized the sharing of de-identified data. Each patient or parent/guardian provided informed consent, as approved by the IRB. To maintain anonymity, all T1 MRI images were defaced prior to publication using an established technique ^67^.

### Subjects

Seventeen subjects (10 female, ages 12 to 43) participated in our study after the placement of 12–17 sEEG electrode leads (Dixi Medical, Marchaux-Chaudefontaine, France) to characterize seizure networks in the context of drug-resistant epilepsy. The clinical epilepsy team planned the targets and trajectories of each sEEG lead based on typical semiology, scalp EEG studies, and brain imaging. No plans were modified to accommodate research. All experiments were performed in the epilepsy monitoring unit or pediatric intensive care unit at the Mayo Clinic in Rochester, MN.

### Data acquisition

Voltage time series were recorded on a g.HighAmp amplifier at 1200 Hz, high-pass filtered above 0.5 Hz in forward and reverse directions with a second-order Butterworth filter, notch filtered at 60, 120, and 180 Hz (± 1 Hz width), and bipolar rereferenced using adjacent channels on the same lead. An electrode located in the white matter far from the hypothesized seizure onset zone(s) was selected as the hardware ground. Data were processed using custom MATLAB code (see Code availability).

### Electrode localization and anatomic labeling

For each subject, the pre-operative T1 MRI was realigned to the anterior and posterior commissure stereotactic space (AC-PC) ^68^, and then coregistered to the post-implant CT using SPM12^69^. Each electrode was subsequently localized using the artefact of each electrode contact seen in the CT scan. Each subject’s T1 MRI was segmented using the Freesurfer 7 autosegmentation algorithm ^70^. Labels were assigned to each vertex of the resulting cortical rendering using the Destrieux cortical atlas ^70^. Bipolar channels were localized as the interpolated position between the two constituent channels. Any channels outside the thalamus and within 2 mm of cortical gray matter were assigned the Destrieux label of the closest cortical vertex. The LeadDBS software ^71^ was used to label thalamic channels according to the Morel atlas ^72^. Specifically, each channel was assigned to all nuclei whose surfaces were within 1 mm such that each bipolar channel could be labeled according to the most common, dominant nuclei between its constituent channels. If multiple nuclei were equally dominant, the bipolar channel received multiple labels.

### Experimental task

All patients performed a roving oddball task that matched the task used in an earlier paper ^7,14^. The task consisted of listening to trains composed of three, five, or eleven identical tones drawn from twenty distinct frequencies (250–6727 Hz, with quarter-octave intervals). These tone trains were presented in a pseudorandom order. Frequencies within each train were identical, but they differed between tone trains, so two consecutive tone trains always had different frequencies. Thus, the first tone of each train was considered deviant because it had a different frequency from the previous tone in the train. Consequently, the last tone of each train was considered a standard because it had the same frequency as the tones that preceded it within that train.

In addition to this standard roving oddball paradigm, an attentional manipulation was added. Subjects therefore heard the same auditory stimuli under two conditions. In the Attended condition, subjects stared at a blank screen and were instructed to focus on the auditory stimuli. In the Unattended condition, subjects performed a concurrent visual counting task: digits were shown on a screen 80 cm away, and subjects counted the occurrences of a prespecified target digit (for example, “3”). The counts reported by each subject were recorded after each Unattended block. Behavioral performance was defined as the proportion of presented target-digit images that were correctly counted. This task was repeated for two unique tone-train sets (oddball wav 1 and 2).

### Event-related potentials

The raw sEEG voltages were transformed into ERP time series as described earlier ^7,14^. In short, bipolar referencing was used to reference the data. To obtain ERPs, the data were down-sampled to 250 Hz and low-pass filtered at 40 Hz. To generate epochs, standard and deviant tones were categorized as described above, and time series from -100 to 500 ms around tone presentation were extracted. Baseline correction was applied by subtracting the mean voltage of the -100 to 0 ms interval from the whole epoch. ERP waveforms were computed as the deviant minus standard difference after baseline correction. Thus, the ERP contrast quantified mismatch-related responses between standards and deviants, but did not by itself isolate prediction error from stimulus-specific adaptation or other state- and history-dependent response components. Electrode-level peak ERP summaries were extracted from the 0–500 ms response window as the signed value of the largest absolute deviation from the standard.

### Mutual Information analyses

We employed Gaussian copula mutual information ^32^ to estimate mutual information between stimulus class (standard/deviant) and the sEEG response (ERP) separately for the Attended and Unattended conditions. The GCMI toolbox estimates mutual information after Gaussian copula normalization of the ERP data (for full details of the method see ^32^), and we have previously implemented it to study neural signals in multiple tasks ^8,14,15,73–75^. Modulated electrodes were defined by significant condition-dependent MI modulation.

For each electrode, stimulus-class labels were permuted 1000 times, with the same permutation applied to both conditions, and Attended and Unattended MI time courses were recomputed to generate a null distribution of condition-difference MI time courses. The maximum and minimum of each permuted difference time course were used to define two-sided maxstatistic thresholds (97.5th and 2.5th percentiles), controlling the family-wise error rate (FWER) at 0.05 across the time-point family within each electrode. No additional max-statistic correction was applied across electrodes; instead, electrode-level detections were summarized at the electrode and participant levels, with participant counts used for Bayesian prevalence inference. Electrodes showing significant positive or negative MI modulation between conditions at any time point were selected as modulated electrodes for further analyses.

For regional summaries, electrodes were included if they showed significant MI modulation and were anatomically assigned to the region being analysed. The resulting electrode set was held constant across the Attended and Unattended conditions. For each participant and condition, regional MI time courses were then obtained by averaging MI across the retained electrodes. Peak MI summaries used the maximum MI value in the 0–500 ms response window:

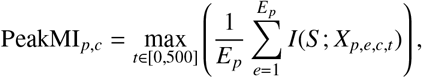

where *p* indexes participants, *c* indexes condition, *E*_*p*_ is the number of retained electrodes for participant *p, S* is stimulus class, and *X*_*p,e,c,t*_ is the ERP value at electrode *e* and time *t*. Temporal participants were assigned to Group A or Group B based on whether their participant-level peak MI was higher in the Attended or Unattended condition.

We computed time-resolved MI across the experiment using overlapping trial windows of 30 trials, advanced in 15-trial steps (50% overlap). For each retained electrode and window, peak MI was extracted as the maximum of the windowed MI time course. Participant-level windowed peak MI was then defined as the mean of these electrode-level peaks,

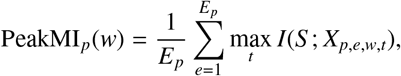

yielding a window-indexed peak-MI time course for each subject and condition. Windowed peak ERP summaries used the same trial windows and retained electrodes, with 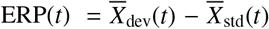 and peak magnitude defined as max_*t*_ | ERP(*t*) | before averaging across electrodes.

### Bayesian Prevalence

It is common practice in cognitive neuroscience to perform null-hypothesis testing, that is, to test whether the results observed in the sample differ from what would be expected if the population average effect were zero. However, participants’ neural responses often differ in many ways, making such inferences difficult. This is especially true for our sample, in which, in addition to individual differences in neurophysiology, the locations of sEEG electrodes were highly heterogeneous across participants.

To overcome this limitation, we employed within-participant (and within-electrode) statistics and reported the number of patients and electrodes that showed significant effects. This approach allows us to employ Bayesian prevalence inference to estimate the prevalence of the observed effects among comparable patients who underwent the same experimental and analytical procedures ^33^. This approach counts the number of patients out of *X* who show an effect and estimates the Bayesian posterior probability of observing the same effect if another comparable patient were randomly selected under the same experimental and analytical procedure.

Bayesian prevalence inference was used to estimate the prevalence of each effect category among comparable patients who underwent the same experimental and analytical procedures, based on the observed number of participants showing a significant within-participant effect (*k*) in the relevant analysis sample (*n*). For each category, the posterior distribution over prevalence was computed and summarized by the posterior median, the maximum a posteriori (MAP) estimate, a one-sided 95% lower-prevalence bound, and 50% and 95% highest-posterior-density intervals (HPDIs). In the temporal analysis, the Bayesian prevalence of Group A and Group B was estimated as mutually exclusive outcome categories within the full temporal-cortex sample (*n* = 17), so patients without significant temporal modulation contributed to the denominator but not to either group-specific numerator. This approach, therefore, converted raw cohort counts into prevalence estimates with quantified uncertainty, rather than reporting sample proportions alone.

### Anatomical control analyses

To assess whether temporal modulation group membership was associated with the anatomical distribution of sampled temporal electrodes, we performed two permutation-based control analyses using the significant temporal electrodes identified in the temporal MI analysis. In the participant-level analysis, participants were assigned to Group A or Group B according to the sign of their participant-level temporal peak MI difference (Attended > Unattended versus Unattended > Attended; peak measured over 0–500 ms). For each participant, significant temporal electrodes were grouped by anatomical label after collapsing hemisphere-specific prefixes (for example, lh_ and rh_), and the number of electrodes in each temporal subregion was counted. These participant-by-subregion counts were summed within Group A and Group B to construct a contingency table. Because the primary null distribution permuted group labels at the participant level while keeping each participant’s full subregion count vector intact, this analysis preserved between-patient differences in the number and anatomical distribution of sampled electrodes.

The observed chi-square statistic was then compared against a null distribution generated by randomly permuting Group A/Group B labels across participants 10,000 times while preserving each participant’s subregion count vector. In a second, pooled electrode-level analysis, all significant temporal electrodes were combined across participants, and each electrode was classified according to the sign of its own peak MI difference (Attended > Unattended or Unattended > Attended). A subregion-by-direction contingency table was constructed, and the observed chi-square statistic was compared against a null distribution generated by randomly permuting electrode-level direction labels 10,000 times.

### Linear mixed-effects modelling

Window-wise participant summaries were analyzed using linear mixed-effects models to compare three candidate temporal profiles: a condition mean-shift model, a linear trend model, and a hinge model with a fixed transition point. For subjects *s* at window *w*, let *y*_*sw*_ denote the outcome measure (PeakMI or PeakERP). A fixed transition point of τ = 7 was used to define a binary post-τ indicator, *C*_*w*_ = *I*(*w* > τ), and a hinge regressor, *H*_*w*_ = max(0, *w* − τ). In two-group analyses, group membership was coded as *G*_*s*_ ∈ {0, 1}. The candidate models were

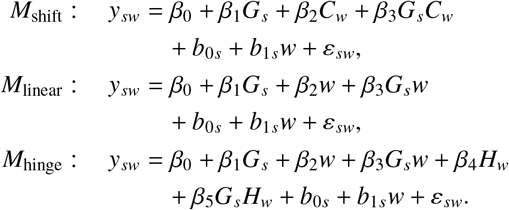

Thus, the mean-shift model tested for an abrupt pre/post-τ change, the linear model tested for a single trend across windows, and the hinge model tested for a change in slope after τ. For one-group analyses, the same models were fitted after omitting the group and interaction terms. Models were fitted by maximum likelihood with subject-specific random intercepts and window slopes; if a model family could not be fitted with the random-slope structure, all candidate models in that comparison were fitted with random intercepts only. Model preference was summarized using log-likelihood and AIC. Because the linear model is nested within the hinge model, this comparison was also evaluated using a likelihood-ratio test and a residual-bootstrap null distribution of the log-likelihood difference. The bootstrap used 1000 resamples with a fixed random seed. In each resample, subject-level residuals from the linear model were resampled within subject, added to the fixed-effect fitted values under the linear null model, and the linear and hinge models were refitted.

The primary temporal grouping separated participants with higher Attended than Unattended peak MI from those with higher Unattended than Attended peak MI, as described above.

As an additional condition-isolated validation of the temporal grouping, we asked whether the original contrast-defined Group A/B labels predicted temporal MI trajectories within each condition separately. For each participant and condition, we used the seven windowed peak-MI values from that condition and estimated a participant-level linear slope,

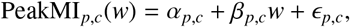

where *w* indexes the condition-local trial window. The group effect was defined as the difference in mean slopes between Group B and Group A. Significance was assessed with an exact permutation test that preserved the original group sizes (*n* = 8 and *n* = 5):all 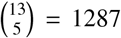 possible assignment of five par-ticipants to Group B and eight participants to Group A were enumerated, and the group-slope difference was recomputed for each assignment. Two-sided *P*-values were calculated as the proportion of permuted absolute group differences greater than or equal to the observed absolute group difference. This procedure was applied separately to the Attended and Unattended trajectories.

Two confirmatory grouping analyses were also used. In the slope-based grouping, participants were assigned based on the sign of their Attended pre-transition temporal MI slope, computed from the first seven Attended windows after centering both window number and peak MI; negative slopes were assigned to Group A and positive slopes to Group B. In the clustering analysis, each participant’s full temporal MI trajectory was standardized within participant and then entered into a deterministic two-cluster k-means algorithm initialized with the two farthest trajectories. Because k-means cluster numbers are arbitrary, the resulting clusters were aligned with the empirical temporal groups based on their average Attended pre-transition MI slope: the lower-slope cluster corresponded to Group A, and the higher-slope cluster corresponded to Group B.

### Co-information analyses

Co-information (co-I) was used to quantify whether prediction-error information at pairs of response time points was redundant or synergistic. For each retained channel, let *X* and *Y* denote the ERP values at two response time points, and let *S* denote the stimulus class. Co-I was defined as

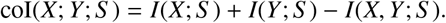

This quantity was evaluated for all pairs of time points, yielding a time-by-time co-I matrix for each channel and condition. Positive co-I values indicate redundancy, where the two time points carry overlapping information about the stimulus class. Negative co-I values indicate synergy, where the joint response across two time points carries information not available from either time point alone.

Empirical co-I was computed separately for each retained temporal channel in the Attended and Unattended conditions, using the same channels selected by the MI-modulated-electrode criterion. Group-level maps were obtained by averaging retained temporal-channel co-I matrices for each participant, then averaging the participant-level maps separately within Group A and Group B. Condition-difference maps were computed as Unattended-minus-Attended. Participant-level condition-difference significance was assessed with stimulus-label shuffles. For each shuffle, standard/deviant labels were permuted within condition, co-I was recomputed for each retained temporal channel, retained channels were averaged within participant, and the Unattended-minus-Attended difference map was recomputed. The maximum and minimum values across the time-by-time matrix from each shuffled participant-level difference map were used to define two-sided max-statistic thresholds, thereby controlling the time-by-time family-within-participant maps. Significance maps counted, at each time-by-time coordinate, the number of participants for whom the observed condition-difference co-I exceeded their participant-specific positive or negative shuffle threshold.

The same co-I analysis was applied to the model data. Model co-I was computed for the first 120 and last 120 simulated trials within temporal layers A1, AB, and PB, restricted to the -100 to 500 ms analysis window. Model group maps were averaged across temporal layers, and model difference maps were computed as Last-minus-First. Difference-map significance was assessed using 250 stimulus-label permutations and max-statistic FWER control, separately for positive and negative effects.

Empirical and simulated co-I difference maps were compared within the 0–500 ms window. Structural similarity (SSIM) quantified similarity in the spatial layout of significant co-I modulation, and signed Pearson correlation quantified whether corresponding significant regions changed in the same or opposite direction. Following the image-based map-comparison approach of Gelens et al. ^14^, significance was assessed using 10,000-permutation null distributions. In each permutation, one map was randomly permuted within the analysis mask, and SSIM or Pearson correlation was recomputed. Permutation *P* values were Bonferroni-corrected across the four planned comparisons: empirical Group A versus simulated Group A, empirical Group B versus simulated Group B, empirical Group A versus empirical Group B, and simulated Group A versus simulated Group B.

### Interpreting MI and co-I results

Importantly, the results of both MI and co-I analyses are reported in bits. As MI can be conceptualized as an effect size of a statistical test of independence ^32^, it is vital to understand how bits differ from more traditional measures of effect sizes, such as Cohen’s D. An increase of one bit corresponds to a halving of the uncertainty about the stimulus class. The effect sizes reported here are the average per sample. Vitally, bits are additive. Thus, since the sampling rate used for our analyses is 250 Hz, an effect size of 0.01 per sample corresponds to 2.5 bits per second. Hence, theoretically, observing the signal for 400 ms with an effect size of 0.01 bits and a sampling rate of 250 Hz should provide 1 bit of information, halving the uncertainty and thus sufficient to identify the stimulus class (standard/deviant).

### Neurocomputational experiments

#### Model architecture and function

To investigate candidate mechanisms underlying the empirical temporal PE responses, we took an existing six-layer-deep neural-network architecture ^39,41^ closely mimicking the neuroanatomy and neurophysiology of six perisylvian areas in the left hemisphere of the human brain involved in spoken language and auditory processing, and adapted it for the present study’s needs.

The prior attention-inhibition simulation motivating this implementation is shown in Supplementary Fig. S2.

This choice was motivated by the fact that the existing neural architecture has been previously used to simulate and explain well-documented neurophysiological patterns of event-related potentials as observed during language processing and oddball stimulation with familiar and unfamiliar sounds ^39,40^.

The model closely reflects the functional and structural features of the mammalian cortex, and incorporates the following neurobiological and neurophysiological constraints:

1. Six cortical areas are modelled, three in the superior temporal and three in the inferior frontal gyri, corresponding to Brodmann Areas (BAs) 41 (A1), 42 (AB), and 22 (PB) in the superior temporal gyrus, and of BAs 44 and 45 (PF), 6V (PM), and 4 (M1) (see Fig. 1**e**);
2. Between-area connections in the model reflect known neuroanatomical links between corresponding brain areas (see next section below); recurrent within-area connections are also modeled, in line with known properties of the mammalian cortex ^76,77^;
3. Between- and within-area links do not implement *all-to-all* connectivity between cells, but sparse, patchy, and topographic projections, with synaptic links established probabilistically (the probability of two cells being connected decreases with the distance; see ^77–79^ and initialized to weak and random efficacy values;
4. Local lateral inhibition ^80,81^ and area-specific global regulation mechanisms (referred to as local and global inhibition, respectively) ^77,81,82^;
5. Single cells’ neurophysiological dynamics, including sigmoid transformation of membrane potentials into neuronal outputs, as well as adaptation and temporal summation of inputs ^83^;
6. Synaptic weights were modified through Hebbian-like long-term potentiation (LTP) and long-term depression (LTD) ^84^, following Garagnani et al. ^39^;
7. Constant presence of uniform uncorrelated white noise (simulating spontaneous baseline neuronal firing) in all model neurons ^85^.

Structurally, each model area consists of two neuronal layers, one of excitatory and one of inhibitory cells, each containing 625 (25×25) cells. Functionally, model cells are graded-response units, each representing a cluster of excitatory pyramidal cells or inhibitory interneurons. The specifics of the computational implementation (including the within-area structure and single-cell functional features) are analogous to those implemented in previously published versions of the architecture (for details, see ^39,40^).

It should be briefly mentioned that our modelling approach, which builds upon and aligns with several previous studies using this neurocomputational architecture ^14,39–42,59,86–88^, was to deploy a fully brain-constrained model. More precisely, we enforced that, for any two model areas, synaptic projections between them are realised only if experimental evidence indicates the presence of neuroanatomical links between the two corresponding cortical areas in the human brain. In the following section, we present the evidence and rationale for the present network architecture.

#### Connectivity of the simulated brain areas

The implemented model areas can be thought of as grouped into two subsystems (frontal and temporal), each simulating a hierarchy of three cortical areas: a primary cortex (motor and auditory, respectively), an adjacent higher secondary, and an associative multimodal region. Neuroanatomical studies in the mammalian brain indicate that adjacent cortical areas tend to be reciprocally connected ^89,90^. We implemented such next-neighbor connections in each of the two subsystems based on known evidence from nonhuman primates: within the frontal / motor (PF–PM–M1) ^89,91^ and within the temporal / auditory (A1–AB–PB) ^92–97^ clusters.

The links connecting the model’s parabelt (area PB) with pre-frontal cortex (PF) are also realised in line with established neuroanatomical evidence on long-distance corticocortical white-matter fibers (arcuate fascicle and extreme capsule) connecting posterior-lateral parts of temporal cortex and inferior prefrontal cortex in humans, and homologue areas in non-human primates ^98–102^.

Finally, the presence of higher-order “jumping” connections between non-adjacent areas in the model (purple arrows in Fig. 1**e**) has been documented by a number of studies, indicating that A1 is directly connected to PB ^89,95–97,103^, that AB is connected to PF ^92,104–107^, that PB and PM are linked ^101,108^ and that PF is also directly connected to M1^109,110^.

#### Procedures

To simulate the roving task, the network was repeatedly presented with stimulus patterns to auditory cortex (area A1). A stimulus pattern, corresponding to an auditory tone, consisted of a predetermined set of 31 cells chosen at random from the 25-by-25 cells of area A1 (about 5% of cells). Twelve different randomly generated stimulus patterns were used; presenting a stimulus involved activating the 31 cells of the selected pattern in A1. A single stimulus presentation consisted of a baseline period with no input, followed by stimulus presentation and an inter-trial interval. A roving sequence was used in which each new deviant was preceded by 6–10 repetitions of the previous standard stimulus, and the new deviant stimulus was chosen at

random. Throughout the simulations, Hebbian synaptic plasticity remained active, strengthening or weakening links between cells whose activities were correlated or anti-correlated.

Simulation outputs comprised 240 trial blocks. Each block contained the recorded activity associated with the final standard response and the subsequent deviant response, sampled at 100 Hz from -300 to 490 ms. For ERP analyses, standard and deviant response epochs were reconstructed from the raw model output, baseline corrected using the prestimulus baseline in the simulated epoch, and converted to deviant-minus-standard responses. Analyses were restricted to the three model areas A1, AB, and PB unless otherwise stated.

The empirical Attended-to-Unattended transition was not simulated directly. Instead, two fixed parameter regimes were analysed as model correlates of different attentional states. Specifically, the high-attention regime (modelling empirical Group A) was implemented by setting the global inhibition parameter (GI) to 90 and the strength of the between-area connectivity (Ffb) to 750. By contrast, the lower-attention regime, simulating empirical Group B, was realised by setting a higher level of global inhibition (GI = 120) and a lower level of conectivity (Ffb = 500).

To match the empirical group sizes, 8 randomly initialised networks were used for Group A and 5 for Group B. For first-versus-last model summaries, the first 120 and last 120 trials were compared within each simulation. Temporal peak MI and maximum positive ERP were extracted from the 0–500 ms response window for each temporal layer and then averaged across A1, AB, and PB. First-versus-last contrasts used paired t-tests across simulations within each model group, with signed-rank tests retained as supplementary non-parametric checks.

For model trajectory analyses, MI and ERP were computed using the same 30-trial windows and 15-trial step used for the empirical windowed analyses. Windows 1–7 came from the first 120 trials and windows 8–14 from the last 120 trials. Model windowed peak MI was defined as the mean across A1, AB, and PB of the maximum MI value in the 0–500 ms response window. Model windowed peak ERP was defined from the deviant-minus-standard model response as the temporal-layer average of the absolute peak response in the reconstructed epoch. Because the model contained first-versus-last segments rather than an explicit task transition, model windowed trajectories were compared with mean-shift and linear mixed-effects models using log-likelihood and AIC.

MI and co-I analyses on the simulated data used the same information-theoretic definitions as the empirical analyses.

## Data availability

This preprint version does not include separate data files.

## Code availability

The MATLAB/Python toolbox GCMI (Gaussian Copula Mutual Information) used in this study is publicly available at: https://github.com/robince/gcmi.

## Acknowledgements

A.C-J. is funded by a Swedish Research Council Project grant (VR; 2025-03245), an ANID/FONDECYT Regular (1240899) and ANID/FONDECYT Regular (1251273) research grants.

## Author contributions

Conceptualization: J.Ä., A.C.-J. Data analysis: J.Ä., M.A.J., M.G. Human sEEG recordings: M.A.J., K.J.M., D.H. Software and methods for human sEEG data: R.I., M.A.J., M.G. Visualization: J.Ä., M.A.J., A.C.-J., M.G. Writing original draft: J.Ä., A.C.-J. Reviewing and editing: T.A.B., M.A.J., M.G., M.V., A.O.B., R.I. Supervision: A.C-J., T.A.B.

## Supplementary figures

**Supplementary Figure S1A.**
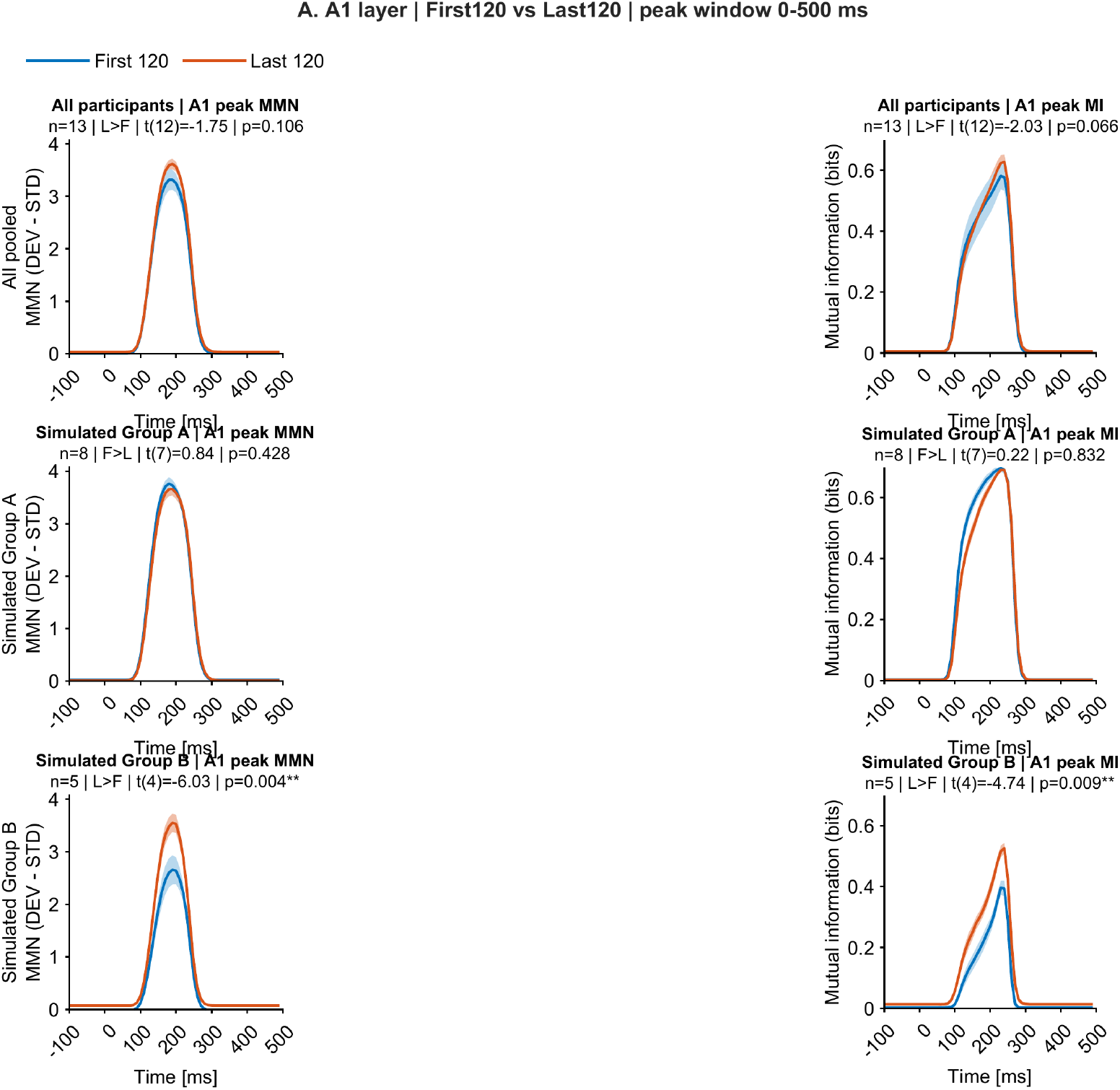
Simulated prediction-error responses in the A1 layer. Left panels show model MMN responses (deviant minus standard), and right panels show stimulus-identity mutual information. First 120 and last 120 trials are compared across all simulations (top), simulated Group A (middle), and simulated Group B (bottom).

**Supplementary Figure S1B.**
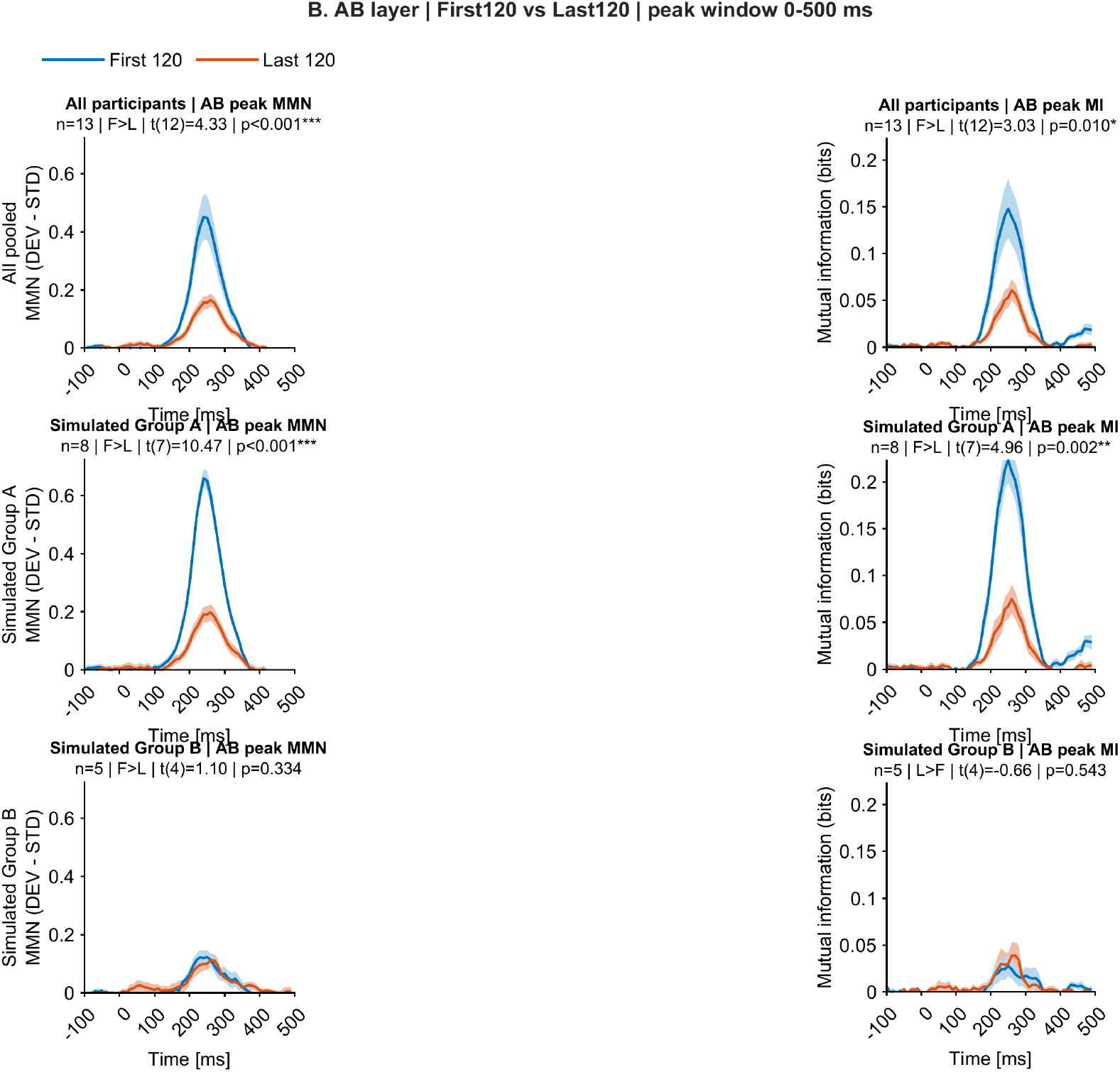
Simulated prediction-error responses in the AB layer. Left panels show model MMN responses (deviant minus standard), and right panels show stimulus-identity mutual information. First 120 and last 120 trials are compared across all simulations (top), simulated Group A (middle), and simulated Group B (bottom).

**Supplementary Figure S1C.**
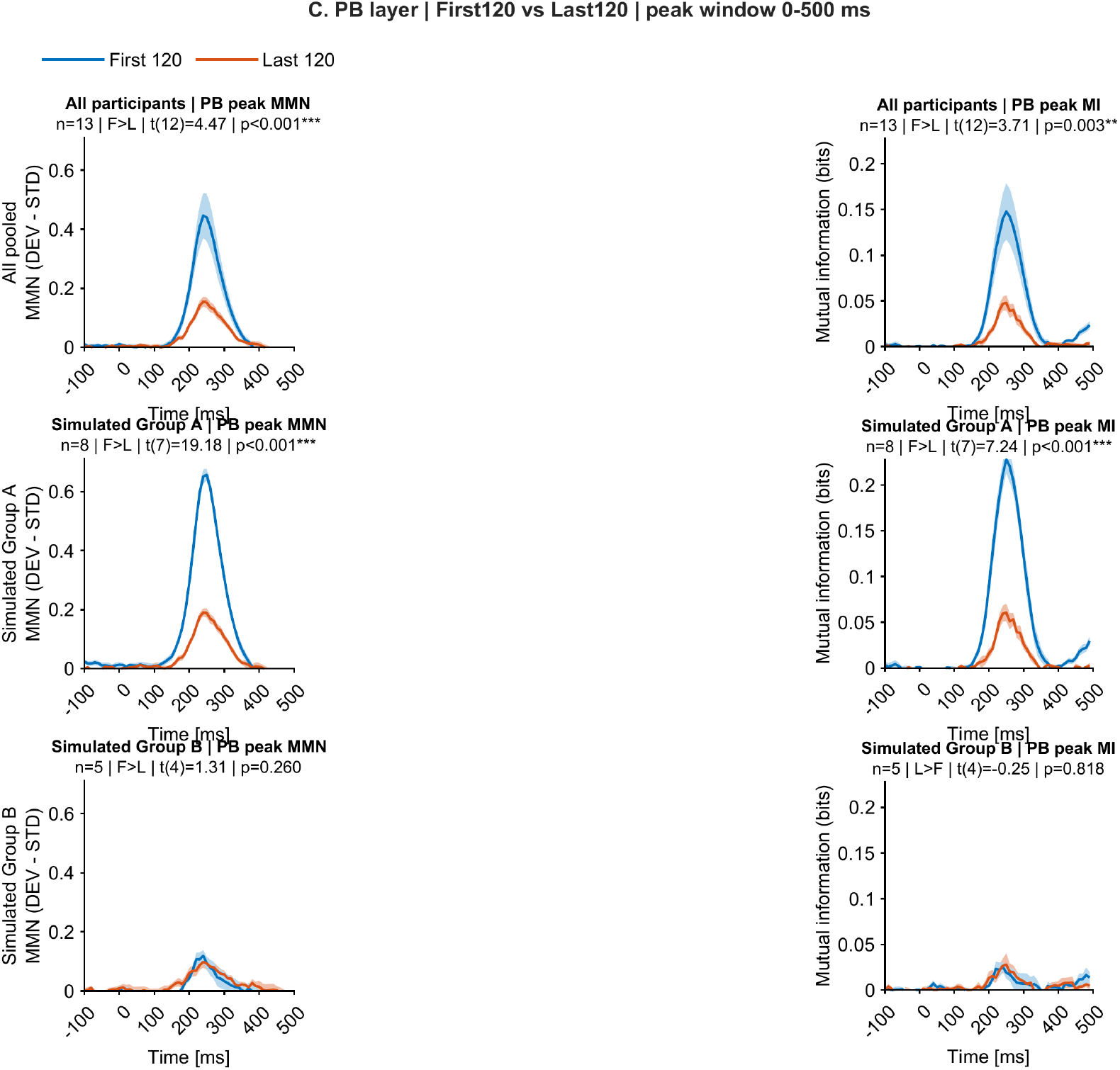
Simulated prediction-error responses in the PB layer. Left panels show model MMN responses (deviant minus standard), and right panels show stimulus-identity mutual information. First 120 and last 120 trials are compared across all simulations (top), simulated Group A (middle), and simulated Group B (bottom).

**Supplementary Figure S2.**
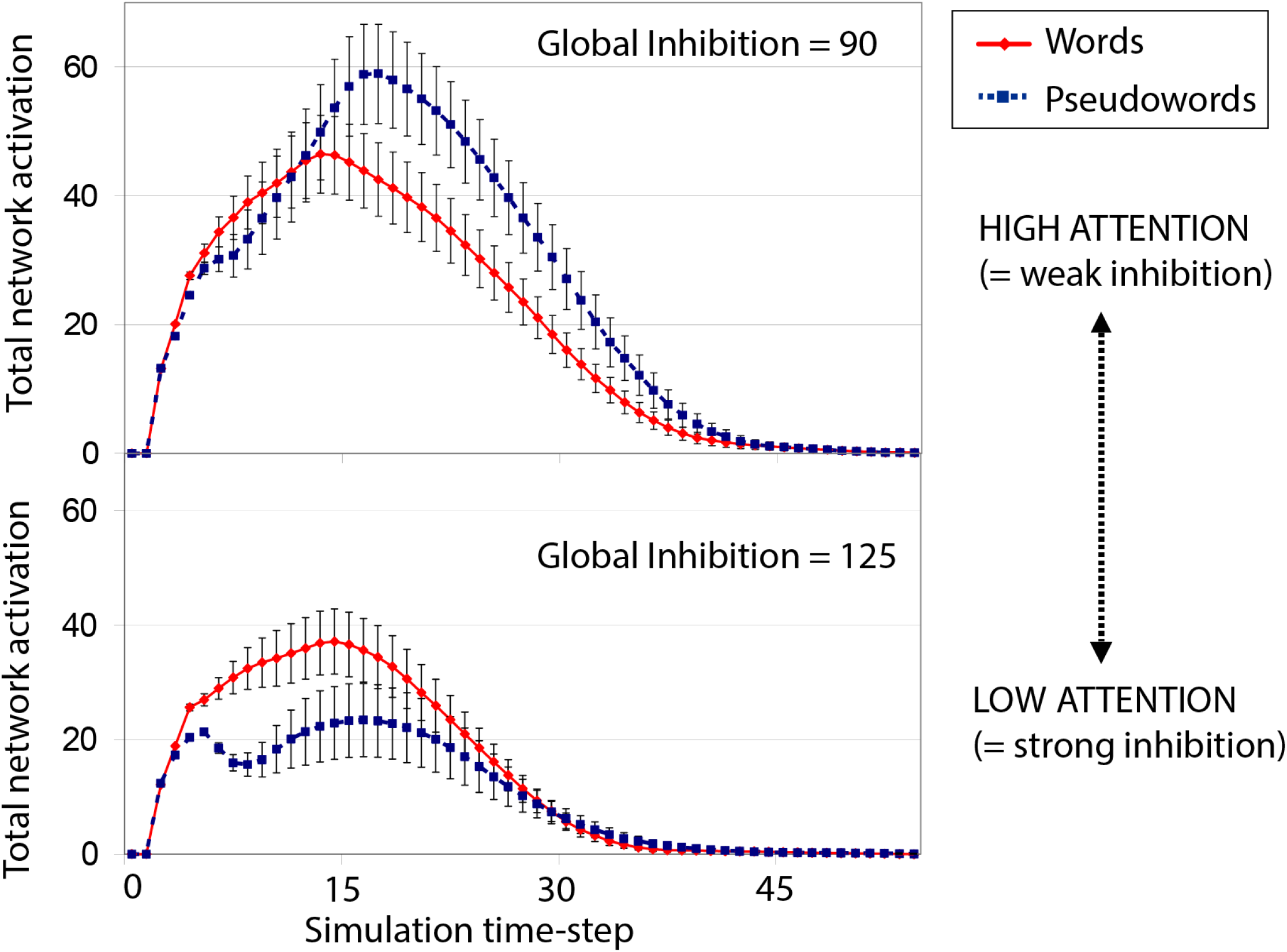
Attention-dependent modulation of familiar and unfamiliar sound responses in the Garagnani neurocomputational architecture. Example simulations from the original model show total network activation for words and pseudowords, corresponding to familiar and unfamiliar auditory patterns, under different global-inhibition regimes. Lower global inhibition (GI = 90) implements a high-attention or weak-inhibition regime, whereas higher global inhibition (GI = 125) implements a low-attention or strong-inhibition regime. The different response profiles across words and pseudowords illustrate the modelling precedent that attentional state can alter the network dynamics evoked by familiar and unfamiliar sounds. In the present simulations, this precedent was extended by pairing GI with between-area connectivity strength (Ffb) to define the Group A and Group B attention regimes. Adapted from ^39^.

